# Temporal scaling of dopamine neuron firing and dopamine release by distinct ion channels shape behavior

**DOI:** 10.1101/2021.08.27.457972

**Authors:** Barbara Juarez, Mi-Seon Kong, Yong S. Jo, Jordan E. Elum, Joshua X. Yee, Scott Ng-Evans, Marcella Cline, Avery C. Hunker, Meagan A. Quinlan, Madison A. Baird, Abigail J. Elerding, Mia Johnson, Derek Ban, Adriana Mendez, Nastacia L. Goodwin, Marta E. Soden, Larry S. Zweifel

**Affiliations:** Department of Psychiatry and Behavioral Sciences, University of Washington; Seattle, WA, United States; Department of Pharmacology, University of Washington; Seattle, WA, United States; Department of Biostructure, University of Washington; Seattle, WA, United States

**Author notes:** University of Maryland, Baltimore; Baltimore, MD. United States. School of Psychology, Korea University; Seoul, Republic of Korea (Yong S. Jo). Icahn School of Medicine at Mount Sinai, New York, NY, United States.

## Abstract

Despite the widely known role of dopamine in reinforcement learning, how the patterns of dopamine release that are critical to the acquisition, performance, and extinction of conditioned responses are generated is poorly resolved. Here, we demonstrate that the coordinated actions of two ion channels, Kv4.3 and BKCa1.1, control the pattern of dopamine neuron firing and dopamine release on different time scales to regulate separate phases of reinforced behavior in mice. Inactivation of Kv4.3 in VTA dopamine neurons increases *ex vivo* pacemaker activity and excitability that is associated with increased *in vivo* ramping dynamics prior to lever press in a learned instrumental response paradigm. Loss of Kv4.3 enhances performance of the learned response and facilitates extinction. In contrast, loss of BKCa1.1 increases burst firing and phasic dopamine release that enhances learning of an instrumental response. Inactivation of BKCa1.1 enhances extinction burst lever pressing in early extinction training that is associated with increased reward prediction error signals. These data demonstrate that temporally distinct patterns of dopamine release are governed by the intrinsic regulators of the cell to shape behavior.

**Teaser:** We show that ion channels in midbrain dopamine neurons are critical for patterning action potential firing at the cell body and governing neurotransmitter release to regulate reinforcement learning.

## Introduction

Reinforcement learning is an essential process by which an organism links environmental stimuli and actions with outcomes. Dopamine-producing neurons in the ventral tegmental area (VTA) of the midbrain facilitate reinforcement learning to promote the performance of goal-oriented behavior through modulation of information processing in the forebrain, most notably in the nucleus accumbens (NAc)(*1*). Distinct patterns of dopamine release that occur on different time scales during the acquisition, performance, and extinction of a conditioned response are thought to underlie dopamine function within these contexts(*2–6*). Despite the evidence that dopamine is critical for reinforcement learning(*1–3*), how the intrinsic properties of the cell contribute to patterning action potential firing and neurotransmitter release during distinct phases of reinforcement learning is not well resolved. Moreover, recent findings suggest that dopamine release dynamics can occur independently of the activity patterns at the cell body(*5*) raising the possibility that the firing properties of dopamine neurons at the level of the cell body may be dispensable for some aspects of reinforcement learning.

VTA dopamine neurons display spontaneous action potential firing in the absence of afferent inputs(*7*) that is regulated by a suite of voltage-gated ion channels (*8–10*). Potassium currents shape the action potential waveform to regulate the frequency and pattern of firing (*8*– *12*). Kv4.3 subunits contribute to A-type potassium currents (*I*_A_) in dopamine neurons that facilitate repolarization and control the frequency of firing (*9*). BKCa1.1 subunits are the essential subunit of voltage-sensitive, calcium-activated big conductance potassium (BK) channels, which conduct BK currents (*I*_BK_). These currents mediate the initial phase of the action potential afterhyperpolarization to control the pattern of firing (*10*). Based on the recruitment of these channels during different phases of the action potential repolarization and afterhyperpolarization, we hypothesized that targeted disruption of their function would differentially impact action potential firing patterns. Specifically, loss of Kv4.3 would induce more pacemaker-like activity and loss of BKCa1.1 would induce more irregular/burst patterns of activity. Thus, inactivation of these channels provides a means to further resolve whether intrinsic regulators of dopamine neuron action potentials influence activity patterns at the level of the cell body and neurotransmitter release dynamics in terminal fields during reinforcement learning.

Using CRISPR/Cas9 gene-editing technologies, we selectively targeted mutagenesis of Kv4.3 (gene name: *Kcnd3*) and BKCa1.1 (gene name: *Kcnma1*) in dopamine releasing neurons of the VTA. We found that inactivation of Kv4.3 and BKCa1.1 differentially affected the action potential waveform, neuronal excitability, and patterns of action potential firing. We further show that loss of function (LOF) of these two channels differentially impacted the acquisition, maintenance, and extinction of a reinforced behavioral response that was associated with distinct effects on the patterns of dopamine release during these behaviors. Specifically, loss of BKCa1.1 enhanced burst firing and phasic dopamine release that was associated with enhanced acquisition of instrumental behavior and a heightened extinction burst. In contrast, loss of Kv4.3 increased dopamine neuron excitability, elevated ramping of dopamine release prior to a lever press in a learned instrumental reinforcement task and facilitated extinction. These findings demonstrate that intrinsic regulators of the action potential waveform influence dopamine release dynamics that control the patterns of behavior during reinforcement learning.

## Results

### CRISPR/Cas9 inactivation of Kv4.3 and BKCa1.1

To determine the distribution of *Kcnd3* and *Kcnma1* expression in the VTA, we performed *in situ* hybridization (RNAscope) to quantify the mRNA levels for these ion channel subunits and the rate-limiting enzyme in dopamine production, *Th*. We find that Kv4.3 and BKCa1.1 subunits are expressed in most dopamine neurons along the rostral-caudal axis of the VTA (Fig. 1, A-C, fig. S1, A-E).

**Fig. 1.**
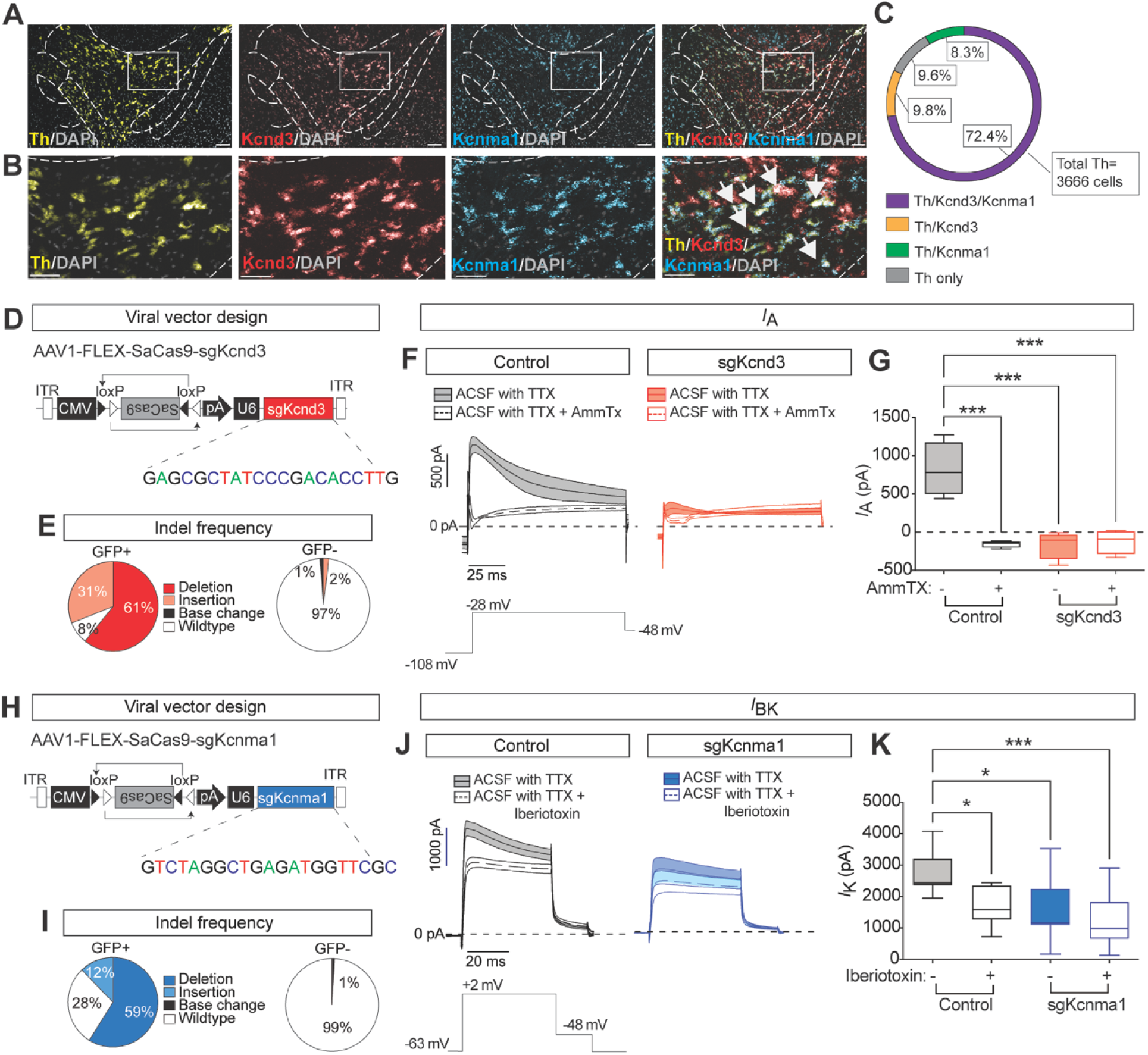
Targeted mutagenesis of *Kcnd3* and *Kcnma1* in dopamine neurons. (**A-B**) RNAscope *in situ* hybridization for *Kcnd3*, *Kcnma1*, and *Th*. White arrows: representative *Th/Kcnd3/Kcnma1* overlap. Scale bar, 100 μm (A) and 50 μm (B). (**C**) Quantification of overlap for *Kcnd3, Kcnma1,* and *Th* in the VTA (*N*=3 mice). (**D**) Schematic of AAV1-FLEX-SaCas9-sgKcnd3 targeting virus. (**E**) Proportion of targeted deep sequencing reads with indel mutations following sgKcnd3 targeting (*N*=5 mice). (**F**) Average pre- and post-AmmTx A-Type currents (*I*_A_) from control (left) and sgKcnd3 (right) -targeted mice (*N*=4 cells/group) and corresponding depolarization step to elicit *I*_A_. (**G**) Average peak *I*_A_ from (**F**) in control and sgKcnd3-targeted mice (Two-way RM ANOVA, Interaction effect F(_1,6_)=18.96, P<0.01; Group effect F(_1,6_)=22.6, P<0.01; Drug effect F(_1,6_)=16.1, P<0.01; Sidak’s Multiple Comparison test, ***p<0.001; *N*_control_=4, *N_s_*_gKcnd3_=4). (**H**) Schematic of AAV1-FLEX-SaCas9-sgKcnma1 targeting virus. (**I**) Proportion of targeted deep sequencing reads with indel mutations following sgKcnma1 targeting (*N*=5 mice). (**J**) Average pre- and post-iberiotoxin potassium currents (*I*_K_) from control (left) and sgKcnma1 (right) -targeted mice (*N*_control_=11 cells; *N*_sgKcnma1_*=*11 cells) and corresponding depolarization step to elicit *I*_K_. (**K**) Peak *I_K_*from (**J**) in control and sgKcnma1 targeted mice following iberiotoxin (Two-way RM ANOVA, Interaction effect F(_1,19_)=18.45, P<0.001; Group effect F(_1,19_)=6.553, P<0.05; Drug effect F(_1,19_)=69.84, P<0.0001; Sidak’s Multiple Comparison test, *p<0.05, ***p<0.001; *N*_control_=11 cells; *N*_sgKcnma1_*=*11 cells). Box and whisker graphs are presented as min-to-max.

To selectively inactivate these two channels, we used a viral-mediated, Cre-inducible CRISPR/SaCas9 system (*13*). Adeno-associated viruses (AAV) containing single guide RNA directed to either of the two loci (AAV1-FLEX-SaCas9-sg*Kcnd3* or AAV1-FLEX-SaCas9-sg*Kcnma1*) were generated (Fig. 1, D and H) and injected into the VTA of separate groups of adult *Slc6a3*(DAT)-IRES-Cre mice (fig. S2, A; fig. S3-S4). Control DAT-IRES-Cre mice received injections of AAV1-FLEX-eYFP into the VTA. We genetically validated gene mutagenesis of targeted sequences by co-injection with AAV-FLEX-KASH-GFP into the VTA and harvesting tissue after four weeks. We found that the AAV mediated Cre-inducible CRISPR/SaCa9 system yielded high efficiency indel formation (Fig. 1 E, I; fig S2 B-C). Next, we used patch-clamp electrophysiology to determine impact of CRISPR/SaCas9 induced Kv4.3 or BKCa1.1 gene mutagenesis on channel activity in VTA dopamine neurons. To visualize cells, we co-injected the LOF mice with AAV-FLEX-eYFP. We found loss of function (LOF) for both channel subunits in dopamine neurons relative to controls (Fig.1, F-G, J-K; fig. S4, A-C). Specifically, AAV1-FLEX-SaCas9-sg*Kcnd3* nearly abolished the AmmTx-sensitive A-type potassium conductance (Fig. 1, F and G) and AAV1-FLEX-SaCas9-sg*Kcnma1* significantly reduced the iberiotoxin-sensitive BK conductance (Fig. 1, J and K; fig. S4, A-C). This loss of sensitivity to channel-specific blockers suggests functional loss of the respective currents.

It is possible that CRISPR/SaCas9-induced LOF of Kv4.3 or BKCa1.1 channels could impact the expression of related potassium channel subunits to compensate for their loss. *Kcnd2* (Kv4.2), another A-type potassium channel subunit, is also expressed in VTA dopamine neurons in mice (*14*). Mutagenesis of *Kcnd3* did not alter midbrain expression of this subunit (fig. S2, D). Previous studies have found no other pore forming alpha subunits for BK channels in the VTA. However, small-conducting calcium activated potassium channels (SK) are expressed in these cells and contribute to the after hyperpolarization (*15*). We found no differences in *Kcnn3* (SK3) expression in the midbrain following *Kcnma1* mutagenesis (fig. S2, E). These data suggest that LOF of either Kv4.3 and BKCa1.1 has significant and specific impacts on their respective currents without inducing compensatory changes in other related channels.

### Kv4.3 and BKCa1.1 regulate *ex vivo* action potential firing but not *ex vivo* terminal release

As predicted, Kv4.3 LOF altered the dopamine neuron action potential waveform by increasing the action potential half width and slowing repolarization relative to control and BKCa1.1 LOF cells (Fig. 2, A-D, control mice). BKCa1.1 LOF depolarized afterhyperpolarization relative to control and Kv4.3 cells and resulted in a more depolarized resting membrane potential relative to control cells (Fig. 2, A-B, E-F). Both Kv4.3 LOF and BKCa1.1 LOF in VTA dopamine neurons significantly depolarized the action potential threshold when compared to control neurons, but neither manipulation affected the peak of the action potential (fig. S4, H-I). To determine if the changes in action potential shape we observed in Kv4.3 LOF and BKCa1.1 LOF dopamine neurons at baseline was impacted by the depolarized resting membrane potential, we clamped the neurons at -60 mV and injected current to induce action potential firing. We found similar effects on action potential shape, although notably no significant change in the anti-peak of the afterhyperpolarization in BKCa1.1 LOF neurons was observed, though the shape of the afterhyperpolarization was affected (fig S4, J-N).

**Fig. 2.**
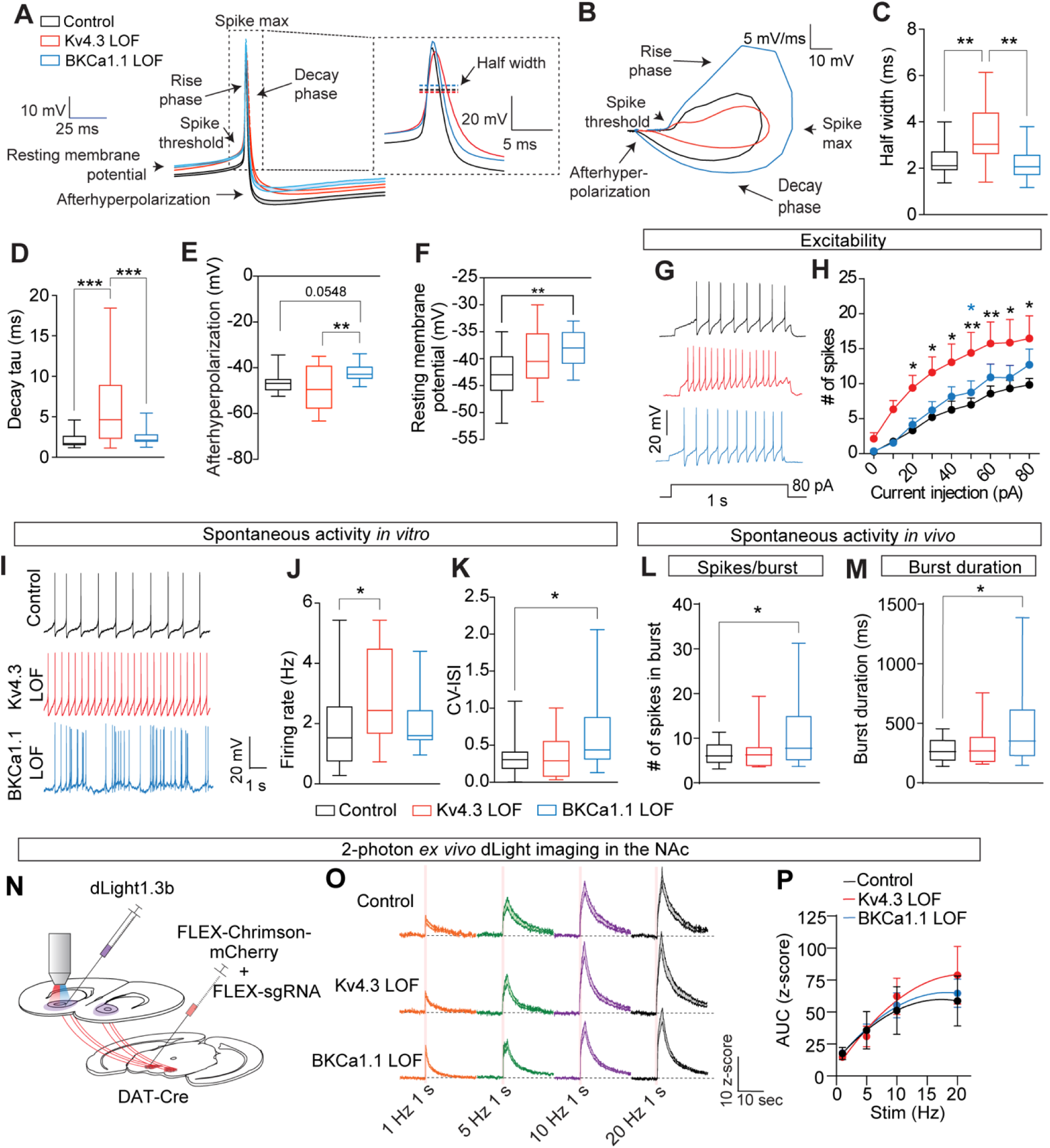
Kv4.3 and BKCa1.1 LOF alter the dopamine action potential waveform and activity. (**A**) Average action potential waveform from spontaneously active VTA dopamine neurons. (**B**) Representative phase plane plot of action potential dynamics. (**C**) Action potential half width in control, Kv4.3 LOF, and BKCa1.1 LOF cells (One-Way ANOVA: F(_2,53_)=9.109, P<0.001, Tukey’s Multiple Comparison’s test **p<0.01). (**D**) Decay kinetics of the action potential (One Way ANOVA: F(_2,53_)=10.67, P<0.0001, Tukey’s Multiple Comparison’s test ***p<0.001). (**E**) Afterhyperpolarization of the action potential (One-Way ANOVA: F(_2,53_)=6.087, P<0.01, Tukey’s Multiple Comparison’s test **p<0.01). (**F**) Resting membrane potential (One-Way ANOVA: F(_2,53_)=6.026, P<0.01, Tukey’s Multiple Comparison’s test **p<0.01). (**G**) Representative evoked excitability traces. (**H**) Current-voltage plot (Two-Way ANOVA, Interaction effect: F(_16, 408_) = 1.143, P=0.3126; Group effect: F(_2, 51_) =4.495 P<0.05; Current effect: F(_8, 408_) =69.36 P<0.0001; Tukey’s multiple comparison’s test: *p<0.05, **p<0.01: black asterisks= Kv4.3 LOF vs control, blue asterisks= Kv4.3 vs BKCa1.1 LOF; *N*_control_=19 cells, *N*_Kv4.3 LOF_=15 cells, *N*_BKCa1.1LOF_=20 cells). (**I**) Representative spontaneous activity traces. (**J**) Firing rate of spontaneous activity (One-Way ANOVA: F(_2,53_)=3.379, P<0.05, Tukey’s Multiple Comparison’s test *p<0.05). (**K**) CV-ISI of spontaneous activity (One-Way ANOVA: F(_2,53_)=3.606, P<0.05, Tukey’s Multiple Comparison’s test *p<0.05). (**C-F** and **J-K**; *N*_control_=21 cells, *N*_Kv4.3LOF_=16 cells, *N*_BKCa1.1_=19 cells). (**L**) Spikes per burst in VTA dopamine neurons *in vivo* (One-Way ANOVA: F(_2,61_)=3.882, P<0.05, Tukey’s Multiple Comparison’s test *p<0.05). (**M**) Burst duration in VTA dopamine neurons *in vivo* (One-Way ANOVA: F(_2,61_)=4.055, P<0.05, Tukey’s Multiple Comparison’s test *p<0.05). (**L-M** *N*_control_=25 cells, *N*_Kv4.3 LOF_=13 cells, and *N*_BKCa1.1_=26 cells). (**N**) Schematic of 2-photon slice imaging of dLight1.3b with optically evoked dopamine release of VTA terminals expressing Chrimson + sgRNA in the NAc. (**O**) Average dLight1.3b signal following 1 s stimulation at different frequencies in control, Kv4.3 LOF and BKCa1.1 LOF mice. (**P**) Average AUC for responses for data presented in panel **O** (Mixed effects analysis, Interaction effect F(_6, 77_)=0.6984, P=0.6516, Group effect F(_2, 26_)=0.05105, P=0.9503, Frequency stimulation effect F_(1.725, 44.28)_=25.99, P<0.0001). (**P-O**, *N*_control_=6 slices; *N*_Kv4.3LOF_=8 slices; *N*_BKCa1.1LOF_=15 slices, except for 5 Hz where *N*_Kv4.3LOF_=7 slices because of brief interruption in data capture that was not detected until post-hoc analysis). Trace averages are presented as mean + SEM. Box and whisker graphs are presented as min-to-max.

Given the changes in action potential waveform dynamics associated with Kv4.3 and BKCa1.1 LOF, we sought to establish whether these perturbations affect neuronal excitability and patterns of action potential firing. Input-output analysis of action potential firing following depolarizing current injection revealed that Kv4.3 LOF in VTA dopamine neurons increased neuronal excitability (Fig. 2, G-H, S4, O) and the frequency of spontaneous action potential firing relative to control cells (Fig. 2 I-J, fig. S4, P). In contrast, BKCa1.1 LOF did not affect these parameters (Fig. 2 G-J, S4, P) but shifted the pacemaker-like *ex vivo* activity observed in control and Kv4.3 LOF cells to a more irregular pattern, as evidenced by an increase in the coefficient of variation in the interspike interval (CV-ISI) (Fig. 2, I and K; fig. S4, P).

Increased CV-ISI in VTA dopamine neurons of BKCa1.1 LOF mice is consistent with a shift to more burst-like activity (*11*); however, this pattern of firing by dopamine neurons is most accurately assessed by recording neural activity *in vivo*. Using optical isolation (*16*) to identify dopamine neurons during *in vivo* recording in the home cage (fig. S5, A-C), we found that BKCa1.1 LOF increases the number of spikes in a burst and burst duration of VTA dopamine neurons relative to controls (Fig. 2, L-M), with no effects on baseline firing rate or the frequency of burst events (fig. S5, D-E). Kv4.3 LOF did not alter *in vivo* firing properties under these basal home cage conditions (Fig. 2, L-M; fig. S5, D-E).

In addition to regulating action potentials at the cell body, potassium channels also regulate neurotransmitter release (*17, 18*) To address whether Kv4.3 or BKCa1.1 LOF function impacts terminally evoked dopamine release dynamics independent from cell body firing, we injected CRISPR/SaCas9 viruses into the VTA along with the redlight-shifted opsin Chrimson(*19*) and injected the genetically encoded dopamine sensor dLight1.3b (*5*) into the NAc. After at least 4 weeks, we performed *ex vivo* 2-photon slice imaging in acute brain slices containing the NAc of optically evoked dopamine release. This *ex vivo* preparation helped us determine if there are any terminal-specific LOF effects on dopamine release, dissociating it from impacts of cell body LOF (Fig. 2, N). Following optical stimulation (1 s) at different frequencies (1, 5, 10, and 20 Hz), we did not detect differences between control and Kv4.3 LOF or BKCa1.1 LOF mice (Fig. 2, O-P).

### Kv4.3 and BKCa1.1 differentially regulate instrumental behavior

To assess whether Kv4.3 and BKCa1.1 LOF impacts reinforcement learning, we first trained mice in an appetitive Pavlovian conditioning paradigm where a lever extension followed by retraction (10s, CS) signals reward (Rew) delivery (Fig. 3A). We did not observe behavioral differences between Kv4.3 LOF and BKCa1.1 LOF mice relative to controls during the CS presentation or ITI period (Fig. 3, B-C). Learning, as assessed by the conditioned approach (CS head entry rate minus ITI head entry rate) was also not different between groups (Fig. 3, D).

**Fig. 3.**
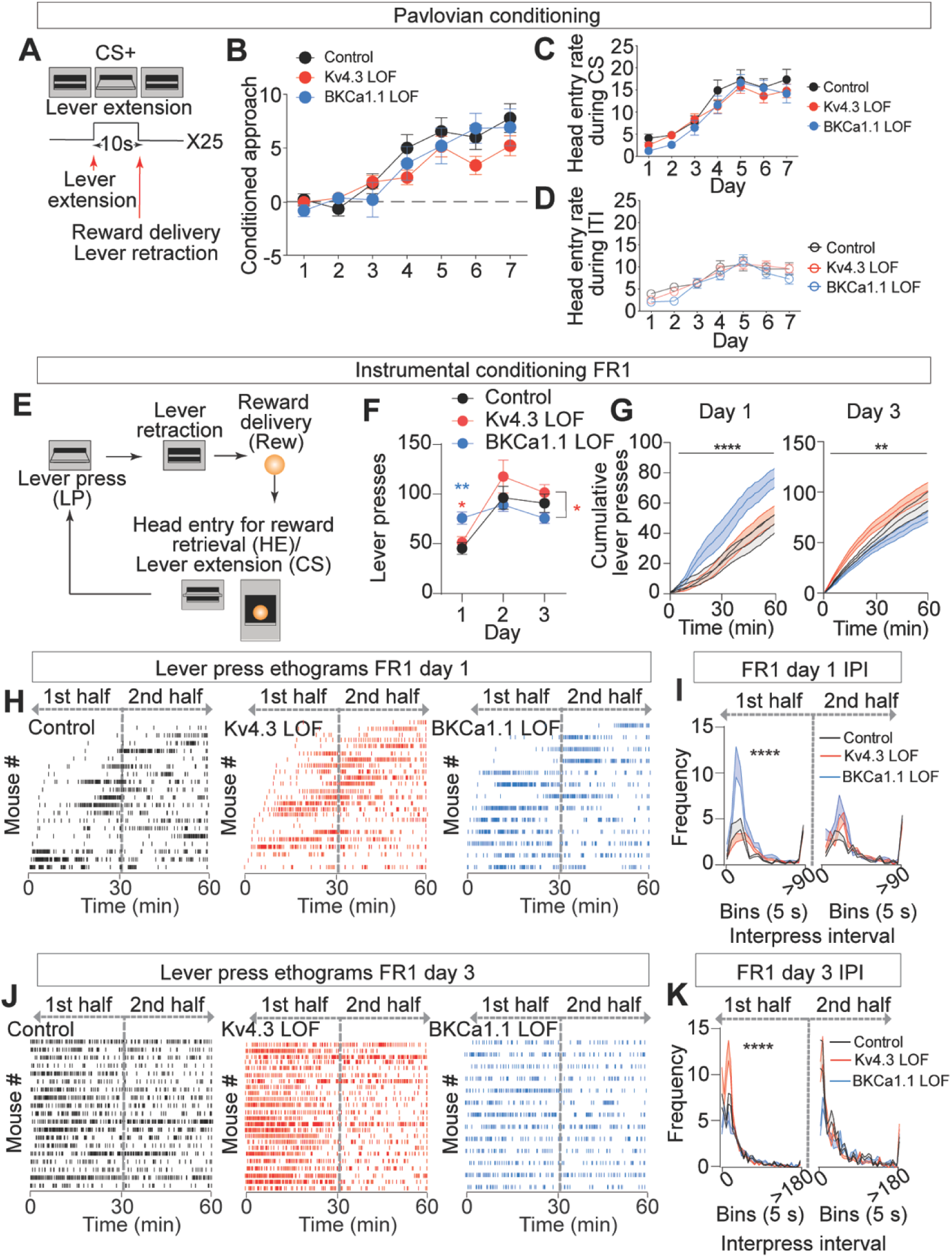
Pavlovian and instrumental behavior in control, Kv4.3, and BKCa1.1 LOF mice. (**A**) Schematic of Pavlovian conditioning paradigm. (**B**) Conditioned approach score (head entries during CS+ - head entries during ITI) across days is not different between groups, (Two-Way RM ANOVA, Interaction effect: F(_12, 378_)=1.557, P=0.1019; Time effect: F(_5, 378_)=31.97, P<0.0001; Group effect: F(_2, 63_)=1.093, P=0.3416) (**C-D**) Head entry rate during CS (C) and ITI period (D) across days used to calculate conditioned approach is not different between groups, (Two-Way RM ANOVA <u>Control ITI vs Cue comparison:</u> Interaction effect F(_6, 276_)=3.86, P<0.01; Time effect F_(3.403, 156.5)_=249, P<0.0001; ITI vs Cue effect F(_1, 46_)=6.433, P<0.05; <u>Kv4.3</u> <u>LOF ITI vs Cue comparison:</u> Interaction effect F(_6, 288_)=2.555, P<0.05; Time effect F_(3.403, 156.5)_=249, P<0.0001; ITI vs Cue effect F(_1, 46_)=6.433, P<0.05; <u>BKCa1.1 LOF ITI vs Cue</u> <u>comparison:</u> Interaction effect F(_6, 204_)=4.286, P<0.0001; Time effect F_(4.053, 137.8)_=36.36, P<0.0001; ITI vs Cue effect F(_1, 34_)=6.989, P<0.05; Sidak’s Multiple Comparison’s test, Cue vs ITI, *p<0.05). (**B-D** *N*_control_=23; *N*_Kv4.3LOF_=25; *N*_BKCa1.1LOF_=18). (**E**) Schematic of instrumental FR1 conditioning paradigm. (**F**) Lever presses during days 1-3 of FR1 conditioning (Two-Way RM ANOVA Interaction effect F(_4, 106_)=2.781, P<0.05; Time effect F_(1.631, 86.42)_=18.26, P<0.0001; Group effect F(_2, 53_)=0.9557, P=0.3911; Subject effect F(_53, 106_)=2.018, P<0.01; Tukey’s Multiple Comparison’s test, *p<0.05, **p<0.01, blue asterisks=BKCa1.1 LOF vs. control; red asterisks= BKCa1.1 LOF vs. Kv4.3 LOF). (**G**) Cumulative lever presses during day 1 (left) and day 3 (right) of FR1 conditioning (Two-Way RM ANOVA, <u>Day 1:</u> Interaction effect F(_118, 3127_)=3.769, ****P<0.0001; Time effect F_(1.807, 95.78)_=143.0, P<0.0001; Group effect F(_2, 53_)=4.832, P<0.05; Subject effect F(_53, 3127_)=167.2, P<0.0001; Tukey’s Multiple Comparison’s test, Kv4.3LOF vs BKCa1.1: p<0.05 for mins 12-17, 19, 40-60; Control vs BKCa1.1LOF: p<0.05 for mins 37-43; p<0.01 for mins 44-60; <u>Day 3:</u> Interaction effect F(_118, 3127_)=1.455, **P<0.01; Time effect F_(1.288,68.26)_=228.8, P<0.0001; Group effect F(_2, 53_)=4.299, P<0.05; Subject effect F(_53, 3127_)=152.3, P<0.0001; Tukey’s Multiple Comparison’s test, Control vs Kv4.3LOF: p<0.05 for mins 2-3, 6-10, 13-20; p<0.01 for mins 4-5; Kv4.3LOFl vs BKCa1.1LOF: p<0.05 for mins 31-37, 42-60; p<0.01 for mins 2-6, 9-18, 20-30, 38-41; p<0.001 for mins 7-8, 19). (**H**) Lever press ethograms during day 1 of FR1 conditioning. (**I**) Frequency distribution of lever inter-press intervals during the first and second half of the conditioning session for day 1 (Two-Way RM ANOVA, <u>first half:</u> Interaction effect F(_72, 1908_)=2.706, ****P<0.0001; Time effect F_(2.888, 153.1)_=20.67, P<0.0001; Group effect F(_2, 53_)=3.458, P<0.05; Subject effect F(_53, 1908_)=3.834; Tukey’s Multiple Comparison’s test, 20 sec: p<0.05 for control vs Kv4.3LOF, Kv4.3LOF vs BKCa1.1LOF; 25 sec: p<0.05 for control vs BKCa1.1LOF; 30 sec: p<0.05 for control vs Kv4.3LOF, p<0.01 for control vs BKCa1.1; 70 sec: p<0.05 for control vs BKCa1.1LOF; 170 sec: p<0.05 for control vs Kv4.3LOF; <u>second half:</u> Interaction effect F(_72, 1908_)=1.012, P=0.4510; Time effect F_(4.049,214.6)_=21.56, P<0.0001; Group effect F_(2, 53)_=1.529, P<0.2263; Subject effect F_(53, 1908)_=2.389, P<0.0001). (**J**) Lever press ethograms during day 3 of FR1 conditioning. (**K**) Frequency distribution of lever inter-press intervals during the first and second half of the conditioning session for day 3 day 3 (Two-Way RM ANOVA, <u>first half:</u> Interaction effect F_(72, 1908)_=1.923, P<0.0001; Time effect F_(3.798, 201.3)_=29.10, P<0.0001; Group effect F_(2, 53)_=4.482, P<0.05; Subject effect F_(53, 1908)_=1.470, P<0.05, Tukey’s Multiple Comparison’s test, 20 sec: p<.0.05 Kv4.3LOF vs BKCa1.1LOF; 25 sec: p<0.01 for Kv4.3LOF vs BKCa1.1LOF; 125 sec: p<0.05 control vs Kv4.3LOF; <u>Second half:</u> Interaction effect F_(72, 1908)_=1.037, P=0.3950; Time effect F_(2.581,136.8)_=19.91, P<0.0001; Group effect F_(2, 53)_=0.6340, P<0.5344; Subject effect F_(53, 1908)_=3.668, P<0.0001). (**F-I:** *N*_control_=19; *N*_Kv4.3LOF_=24; *N*_BKCa1.1LOF_=13). **B-D** and **F** data are presented as mean ± SEM.Cumulative lever press trace data presented as mean ± SEM and IPI data presented as mean + SEM.

Following Pavlovian conditioning, mice were transitioned to a fixed-ratio (FR1) schedule of operant reinforcement learning in which lever retraction and reward delivery were contingent on a lever press (LP). Extension of the lever that had previously served as the onset of the CS was now contingent on making a head entry (HE) into the food hopper (Fig. 3, E); thus, serving as a conditioned reinforcer. BKCa1.1 LOF learned the contingency faster than control and Kv4.3 LOF mice on day one of training, (Fig. 3, F-G). Control and Kv4.3 LOF mice increased responding between the first and second day that remained elevated on the third day, with no differences in total lever presses between groups (Fig. 3, F). However, Kv4.3 LOF mice had a higher initial rate of responding than control or BKCa1.1 mice on the third day, once the task was acquired (Fig. 3, G). Analysis of lever press ethograms revealed different patterns of operant responding between the groups (Fig. 3, H-J). Consistent with the faster learning of the contingency, BKCa1.1 LOF mice started pressing more quickly and with a shorter interpress interval on the first day (Fig. 3, H-I). Once the contingency was learned, control and BKCa1.1 LOF mice had a relatively stable level of responding on the third day; however, Kv4.3 LOF mice showed a high level of responding in the first half of the session that diminished in the second half (Fig. 3, J-K).

Differences in reinforcement learning were not associated with changes in overall locomotor activity (fig. S6, A-B). To assess whether the differences in operant responding reflected altered motivation, we assayed mice in a progressive ratio schedule of reinforcement. We did not observe differences between the groups in this task (fig. S6, C).

### Kv4.3 and BKCa1.1 differentially regulate behaviorally-evoked firing patterns and dopamine release

Differential regulation of action potential waveforms, intrinsic excitability, and patterns of instrumental behavior suggest that Kv4.3 and BKCa1.1 LOF cause distinct changes in the encoding of reward-related behaviors. To test this, we performed *in vivo* electrophysiology during the instrumental conditioning task with optical identification of VTA dopamine neurons (Fig. 4, A,B) following Pavlovian conditioning as above. Due to the restrictive nature of the tethering for recording, mice underwent 5 days of FR1 instead of 3 days and cell body activity was measured on day 1 and day 5. On the first day of conditioning, we detected 13 optically sensitive units in controls, 15 optically sensitive units in Kv4.3 LOF, and 15 optically sensitive units in BKCa1.1 mice. Of these, 7 control, 14 Kv4.3, and 10 BKCa1.1 optically sensitive dopamine neurons were activated by the lever press (LP) and reward delivery (Rew) (Fig. 4, C-E). On the fifth day of conditioning, we detected 7 optically sensitive units in controls, 13 in Kv4.3 LOF, and 19 in BKCa1.1 LOF mice. 7 of 7 cells were activated in controls, 9 of 13 in Kv4.3 LOF, and 13 of 19 in BKCa1.1. Consistent with the increased ‘burstiness’ of dopamine neurons in BKCa1.1 LOF mice, we observed higher responses to the LP in optically sensitive cells in BKCa1.1 LOF mice on the first day of conditioning relative to control and Kv4.3 LOF mice (Fig. 4, D) and on the fifth day of conditioning in BKCa1.1 LOF mice relative to Kv4.3 LOF mice (Fig. 4, D). Although Rew responses were generally higher in BKCa1.1 LOF mice, we only observed significance on the fifth day of conditioning between BKCa.1.1 LOF and Kv4.3 LOF mice (Fig. 4, E).

**Figure 4.**
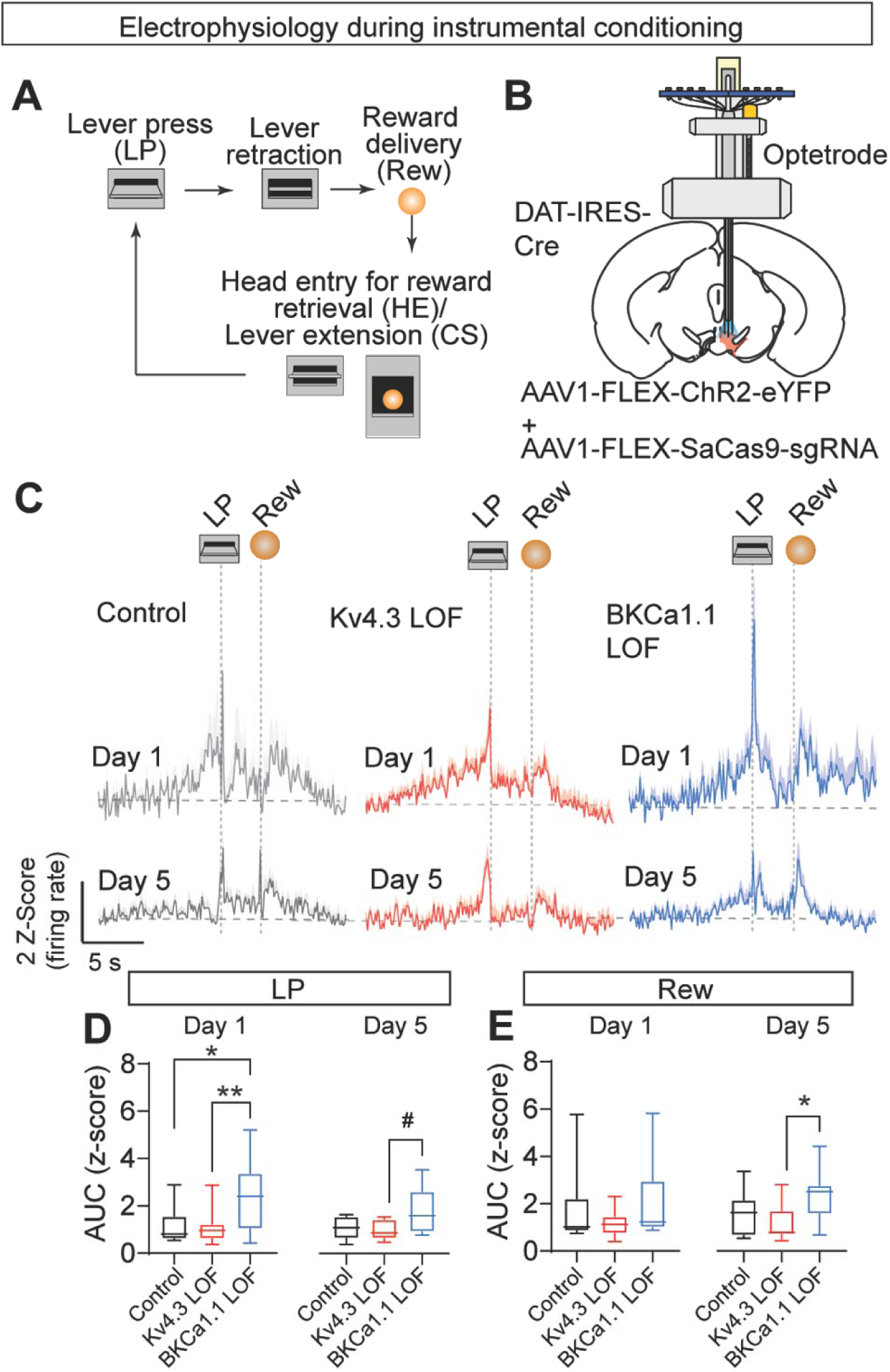
Differential action potential firing in dopamine neuron from control, Kv4.3 LOF, and BKCa1.1 LOF mice during instrumental conditioning. (**A**) Schematic of instrumental conditioning paradigm. (**B**) Illustration of optetrode Microdrive placement in the VTA and injections of AAV1-FLEX-ChR2-eYFP + AAV1-FLEX-SaCas9-sgRNA injection in DAT-IRES-Cre mice. (**C**) Average peri-event Z-score for optically sensitive dopamine neurons responsive to LP in the VTA for day 1 (top row) and day 5 (middle row and bottom row) for (control (*N*=2), Kv4.3 LOF (*N*=4), and BKCa1.1 LOF (*N*=3) mice. (**D**) Average area under the curve (AUC) for the peri-event Z-score action potential firing post-LP period (0.7 s) on day 1 (One-way ANOVA, F_(2,28)_= 4.62, P=0.014, Tukey’s multiple comparisons, *p<0.05, **p<0.01; *N*_control_=7; *N*_Kv4.3LOF_=14; *N*_BKCa1.1LOF_=10) and day 5 (One-way ANOVA, F_(2,26)_= 4.067, P=0.029, Tukey’s multiple comparisons, #p=0.0502, *N*_control_=7; *N*_Kv4.3LOF_=13; *N*_BKCa1.1LOF_=19). (**E**) Average AUC for the Z-score of action potential firing for post-Rew period (2 s) on Day 1 and Day 5 (One-way ANOVA, F_(2,26)_= 3.94, P=0.032, Tukey’s multiple comparisons, *p<0.05; Day 1: *N*_control_=7; *N*_Kv4.3LOF_=14; *N*_BKCa1.1LOF_=10, Day 5: *N*_control_=7; *N*_Kv4.3LOF_=13; *N*_BKCa1.1LOF_=19). Averaged traces are presented as mean + SEM. Box and whisker graphs are presented as min-to-max.

To determine whether changes in behaviorally-evoked action potential firing at the cell body associated with ion channel LOF is related to dopamine release dynamics downstream, we expressed dLight1.3b in the NAc of Kv4.3 LOF, BKCa1.1 LOF, and control mice (fig. S7) and performed fiber photometry during FR1 following Pavlovian conditioning as above. Again, mice underwent 5 days of FR1 instead of 3 days and dopamine signals were measured on day 1 and day 5. Control mice showed phasic increases in dopamine time-locked to LP and Rew on day 1 (Fig. 5, A top row) and day 5 (Fig. 5, A middle row, bottom row). Kv4.3 LOF mice showed highly variable LP and Rew dopamine signals on the first day (Fig. 5, A top row); clear signals to these events emerged on day 5 (Fig. 5, A middle row, bottom row heat maps). BKCa1.1 LOF mice had large dopamine release events to LP and Rew on day 1 (Fig. 5, A top row) and day 5 (Fig. 5, A middle row, bottom row). Similar to cell body firing BKCa1.1 LOF mice had larger dLight1.3b signals during LP on day 1 relative to Kv4.3 LOF mice and showed a trend towards higher signals on day 5 relative to control mice (P=0.074) (Fig. 5, B). Rew-evoked signals were also significantly higher in BKCa1.1 LOF mice on day 1 compared to control and Kv4.3 LOF mice (Fig. 5, C). As with action potential firing, we observed prominent ramps in the dopamine signal prior to LP on day 1 that largely diminished in control mice by day 5 (Fig. 4, C; Fig. 5, A, D-G). In Kv4.3 LOF mice this ramp remained prominent on day 5 in both the dLight1.3b fiber photometry and optetrode recordings that was significantly higher than controls (Fig. 5, D-G).

**Figure 5.**
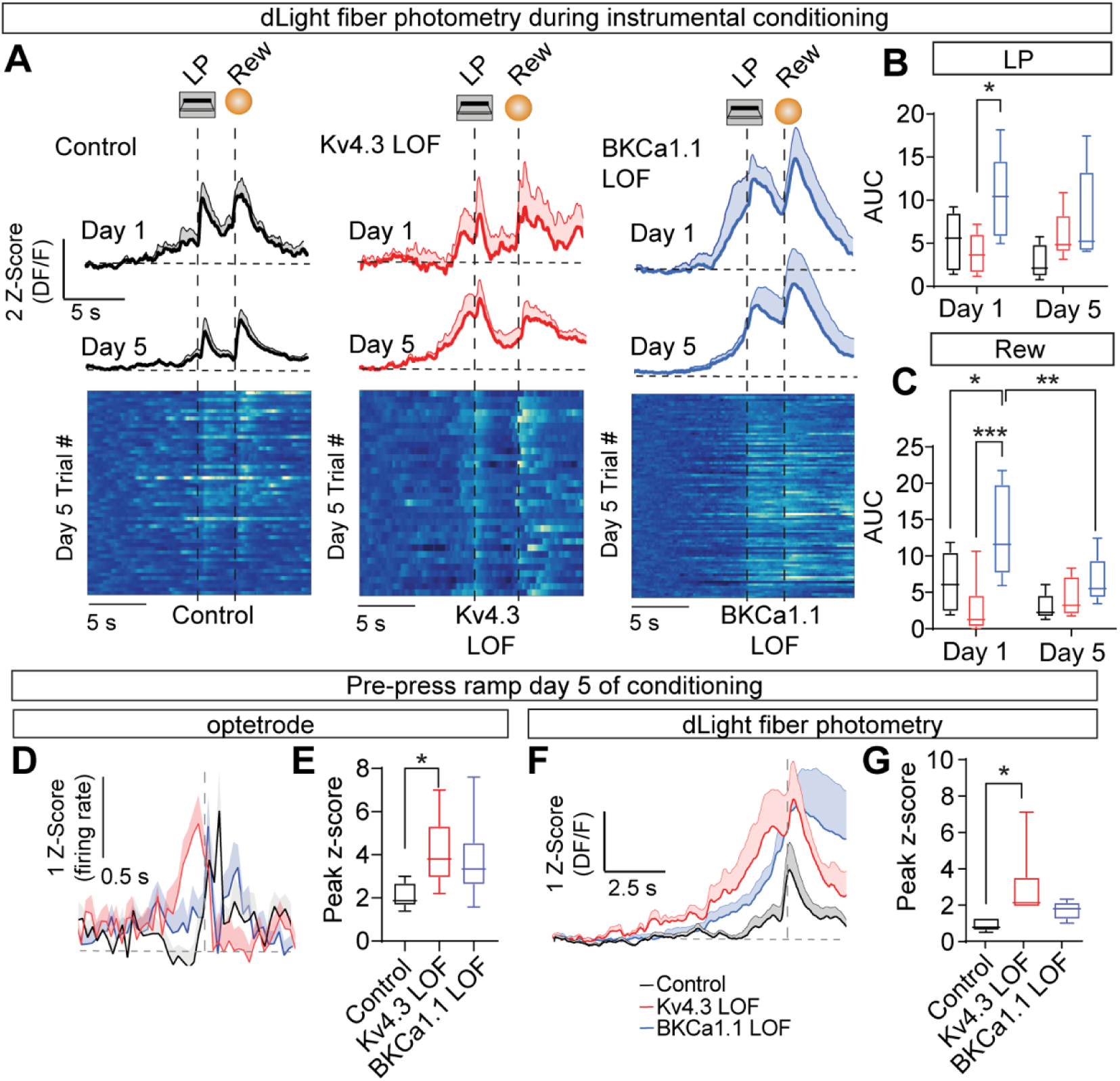
Differential dopamine dynamics in the NAc of control, Kv4.3 LOF, and BKCa1.1 LOF mice during instrumental conditioning. (**A**) Average peri-event Z-score for dLight1.3b signals in the NAc for day 1 (top row) and day 5 (middle row and bottom row) for (control (*N*=6), Kv4.3 LOF (*N*=6), and BKCa1.1 LOF (*N*=5) mice. (**B**) Average AUC for the perievent Z-score dLight1.3b signals post-LP period (3 s) (Two-Way RM ANOVA Interaction effect F_(2, 14)_ =1.564, P=0.24;Day effect F_(1, 14)_=0.556, P=0.47; Group effect F_(2, 14)_=5.472, P<0.05; Subject effect F_(14, 14)_=1.156, P=0.39, Sidak’s Multiple Comparison’s test *p<0.05). (**C**) Average AUC for the Z-score for post-Rew period (5 s) (Two-Way RM ANOVA Interaction effect F_(2, 14)_=6.766, P<0.01;Day effect F_(1, 14)_=9.774, P<0.01; Group effect F_(2, 14)_=5.361, P<0.05; Subject effect F_(14, 14)_=3.293, P<0.05, Tukey’s or Sidak’s Multiple Comparison’s test, *p<0.05, **p<0.01, ***p<0.001). (**D**) Zoomed average peri-event Z-score on day 5 for optically-sensitive neurons in the VTA from 4C showing ramping in the pre-LP period in control (*N*=2), KV4.3 LOF (*N*=4), and BKCa1.1 LOF (*N*=3) mice. (**E**) Average ramp to peak Z-score from optically sensitive neurons in the VTA during the pre-LP period (One-way ANOVA, F_(2,25)_= 1.944, P=0.029, Tukey’s multiple comparisons, *p<0.05). (**F**) Zoomed average peri-event Z-score on day 5 for dLight1.3b in the NAc from panel A showing ramping in the pre-LP period. (**G**) Average ramp to peak Z-score from dLight1.3b signals in the NAc for pre-LP period (One-way ANOVA, F_(2,14)_=3.848, p<0.05, Tukey’s Multiple Comparison’s test *p<0.05). (**A-C, F-G:** *N*_control_=6; *N*_Kv4.3LOF_=6; *N*_BKCa1.1LOF_=5). Averaged traces are presented as mean + SEM. Box and whisker graphs are presented as min-to-max.

### Kv4.3 and BKCa1.1 differentially regulate extinction of an operant response

Following Pavlovian and FR1 conditioning, a subset of mice underwent extinction training with reward omission (Fig. 6, A). Control mice exhibited an extinction burst in lever pressing on the first day of extinction (higher pressing on day 1 of extinction versus the last day of reinforced FR1 conditioning). This extinction burst was elevated in BKCa1.1 LOF mice relative to control mice (Fig. 6, B-C; fig. S8, A-C). In contrast, Kv4.3 LOF mice had significantly reduced lever presses during the first day of extinction training relative to both control and BKCa1.1 LOF mice (Fig. 6, B-C; fig. S8, A-C). The number of lever presses in BKCa1.1 LOF and control mice decreased during the extinction session but remained relatively stable in Kv4.3 LOF mice (Fig. 6, C; fig S8. A-B). All mice performed similarly on the last day of extinction (Fig. 6, B; fig. S8, D-F). Interestingly, lever pressing on the first day of extinction was proportional to the level of responding on the first day of FR1 conditioning (fig. S8, G), suggesting that enhanced acquisition is associated with an invigorated response when the outcome (reward omission) does not match expectation (reward).

**Fig. 6.**
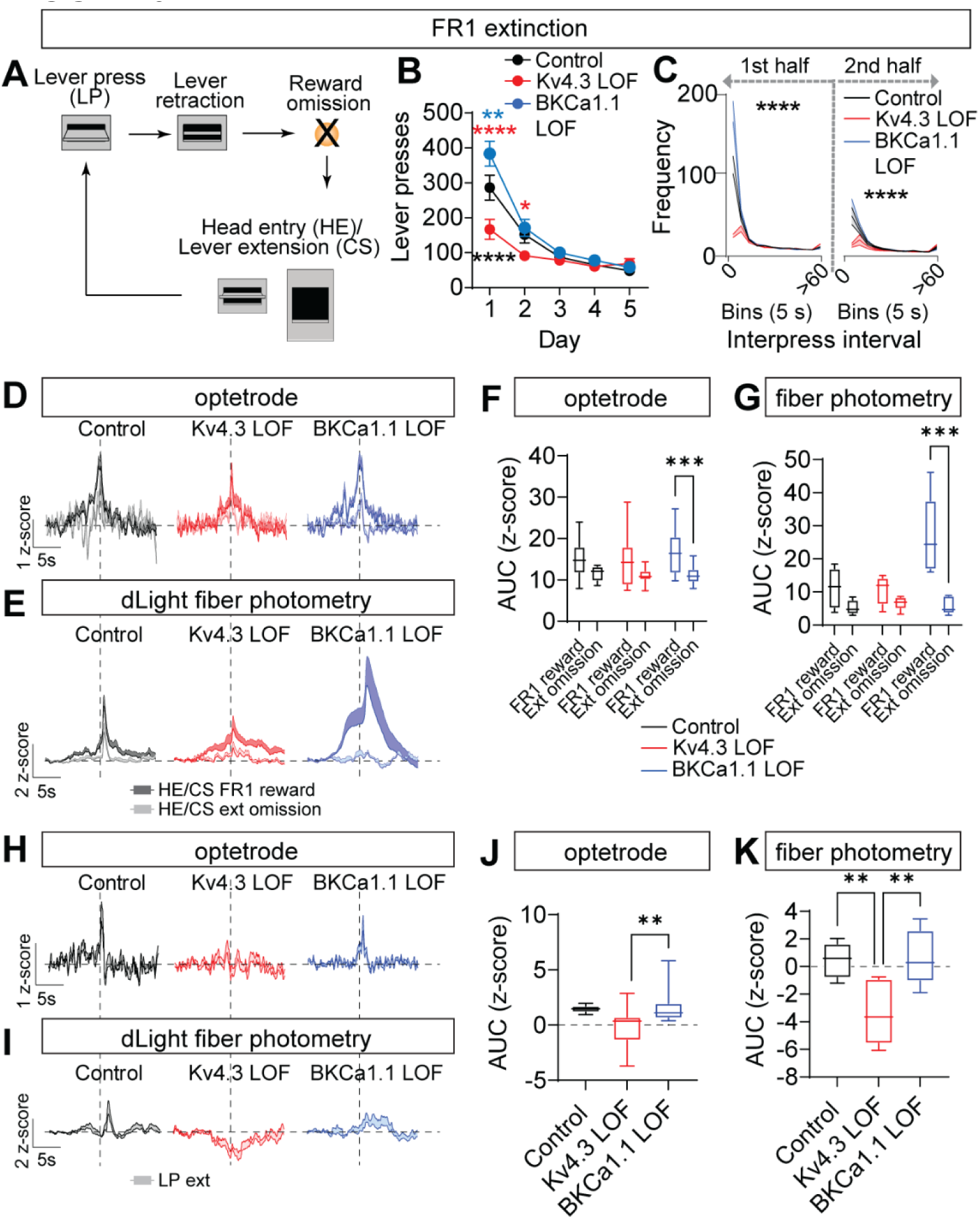
Extinction behavior and dopamine signal diametrically changes in Kv4.3 and BKCa1.1 LOF mice. (**A**) Schematic of instrumental FR1 extinction paradigm. (**B**) Lever presses during days 1-5 of extinction training (Two-way RM ANOVA, Interaction effect F_(8, 128)_=8.674, P<0.0001; Time effect F_(1.785, 57.13)_=79.88, P<0.0001; Group effect F_(2, 32)_=7.744, P<0.001; Subject effect F_(32, 128)_=2.872, P<0.0001; Tukey’s Multiple comparison’s test, **p<0.01, blue= BKCa1.1 LOF vs. control, ****p<0.0001, red=BKCa1.1 LOF vs. Kv4.3 LOF, ****p<0.0001, black= Kv4.3 LOF vs. control). (**C**) Frequency distribution of lever presses during the first and second half of the conditioning session for day 1 of extinction (Two-way RM ANOVA, <u>first half:</u> Interaction effect F_(72, 1152)_=10.31, ****P<0.0001; Time effect F_(1.141, 36.51)_=57.07, P<0.0001; Group effect F_(2,32)_=12.42, P<0.0001; Tukey’s multiple comparison’s test, 5 sec: p<0.01 for control vs Kv4.3LOF and Kv4.3LOF vs BKCa1.1LOF; 10 sec: p< 0.05 for Kv4.3LOF vs BKCa1.1LOF; 25 sec: p<0.05 for Kv4.3LOF vs BKCa1.1LOF; 45 sec: p<0.05 for control vs BKCa1.1LOF; 60 sec: p<0.05 for Kv4.3LOF vs BKCa1.1LOF; <u>second half:</u> Interaction effect F_(72, 1152)_=5.047, P<0.0001; Time effect F_(1.459, 46.60)_=48.96, P<0.0001; Group effect F_(2, 32)_=5.851, P<0.01, Tukey’s multiple comparison’s test, 5 sec: p<0.05 for control vs Kv4.3LOF, p <0.01 for Kv4.3LOF vs BKCa1.1LOF). (**B-C**, *N*_control_=13, *N*_Kv4.3LOF_=10, and *N*_BKCa1.1LOF_=12;) (**D**) Average Z-score for optically-sensitive dopamine neurons in the VTA during day 1 and day 5 following rewarded (FR1 reward) HE/CS compared to average Z-score following non-rewarded HE (ext omission) during extinction in control (*N*=2), KV4.3 LOF (*N*=4), and BKCa1.1 LOF (*N*=3) mice. (**E**) Average Z-score for dLight1.3b signals in the NAc during day 1 and day 5 following FR1 rewarded HE/CS trials compared to average Z-score following non-rewarded Ext omission trials during extinction in control (*N*=6), KV4.3 LOF (*N*=6), and BKCa1.1 LOF (*N*=5) mice for day 1 of extinction. (**F**) Average AUC for the Z-score of action potential firing in the VTA for HE/CS period during FR1 reward trials and Ext omission trials (Two-way repeated measures ANOVA, Group Effect: F_(2, 86)_=0.9786, P=0.38; Time effect F_(1, 86)_=19.55, P<0.0001, Sidak’s within group comparisons, ***p<0.001). (**G**) Average AUC for the Z-score of dLight1.3b signals in the NAc for HE/CS period during FR1 rewarded and non-rewarded Ext omission trials (Two-way repeated measures ANOVA, Interaction effect F_(2, 14)_=8.389, P<0.004; Group effect F_(2, 14)_=5.984, P=0.0132; Time Effect F_(1, 14)_=30.59, P=0.0001; Sidaks’s within group comparisons, ***p<0.001). (**H**) Average Z-score for optically sensitive neurons in the VTA during lever press (LP) period during extinction day 1 in control (*N*=2), KV4.3 LOF (*N*=4), and BKCa1.1 LOF (*N*=3) mice. (**I**) Average Z-score for dLight1.3b signals in the NAc during extinction day 1 in control (*N*=6), KV4.3 LOF (*N*=6), and BKCa1.1 LOF (*N*=5). (**J**) Average AUC for the Z-score of action potential firing in the VTA for lever press period during extinction day 1 (One-way ANOVA, F_(2, 40)_=6.406, P=0.0039; Tukey’s multiple comparisons, **p<0.0036). (**K**) Average AUC for the Z-score of dLight1.3b signals in the NAc for lever press period during extinction day 1 (One-way repeated measures ANOVA, F_(2, 14)_=8.443, P=0.0039; Tukey’s multiple comparisons, **p<0.01). Data are presented as mean ± SEM (**B-E**, **H-I**), or Box and Whisker min-to-max graphs (**F-G**, **J-K**).

Next we sought to determine the impact of reward omission during extinction training on action potential firing and dopamine release. We compared dopamine neuron firing and NAc dopamine release responses in extinction training during the first head entry into the food port (HE)/re-extension of the lever (CS) following a lever press to the average reinforced HE/CS responses during acquisition training (Fig. 6, D-E). Control and Kv4.3 LOF mice did not show a significant difference in dopamine neuron action potential firing and NAc dLight signals during HE/CS period in extinction relative to acquisition (Fig. 6, D-G). In contrast, the dopamine neuron action potential firing and dLight signals of BKCa1.1 LOF during HE/CS events of extinction training with reward omission were significantly reduced when compared to HE/CS events of acquisition training with reward delivery (Fig. 6, D-G).

The dramatic differential responding in dopamine neuron firing and release during HE/CS events in BKCa1.1 LOF mice may explain the elevated extinction burst lever pressing, but this does not explain the reduced lever press responding in Kv4.3 LOF mice. To explore this further, we analyzed dopamine neuron action potential firing and NAc dopamine release during the LP period of extinction training on day 1 (Fig. 6, H-I). Action potential firing and dopamine release to the LP were detectable in control and BKCa1.1 LOF mice, but these signals were significantly reduced in Kv4.3 LOF mice (Fig. 6, H-K). Interestingly, while no differences were observed in dopamine neuron action potential firing in Kv4.3 LOF mice, NAc dLight signals dipped significantly below baseline to the LP (Fig. 6, H-K).

## Discussion

Our results show that the ion channels that contribute to the generation of the action potential regulate the patterns of dopamine neuron firing and dopamine release to influence reinforcement learning, performance, and extinction. Using viral delivery of CRISPR/SaCas9 we were able to generate high-efficiency mutagenesis that allowed us to determine the cell-type specific function of Kv4.3 and BKCa1.1 in dopamine neurons of the adult brain. We found that CRISPR/SaCas9 mutagenesis of *Kcnd3* (Kv4.3 LOF) effectively reduced the AmmTx-sensitive A-type potassium current and mutagenesis of *Kcnma1* (BKCa1.1 LOF) significantly attenuated the iberiotoxin-sensitive BK current. Loss of these channels altered the action potential waveform by broadening the action potential half-width (Kv4.3 LOF) and reducing the AHP (BKCa1.1 LOF). Reduced AHP in VTA dopamine neurons following loss of BKCa1.1 is distinct from the effects of inhibition of BK current in substantia nigra pars compacta (SNc) dopamine neurons where a paradoxical increase in afterhyperpolarization was reported (*10*). This could be due in part to the higher expression of the small conductance potassium channel, SK3 in SNc dopamine neurons (*20*) that also contributes to the afterhyperpolarization (*15*), or the reciprocal interactions between Kv2 channels and BK channels in SNc dopamine neurons (*10*).

Kv4.3 LOF resulted in an increase in VTA dopamine neuron excitability and pacemaker activity, as would be predicted based on the recruitment of A-type potassium channel conductance during the rapid depolarization phase of the action potential to generate rapid feedback and allow for slow pacemaker activity (*9*). Loss of BKCa1.1 resulted in irregular action potential firing in VTA dopamine neurons *ex vivo* and prolonged burst events in these cells *in vivo*. This again contrasts to the effects observed in SNc dopamine neurons, likely relating to the differential effects on the AHP. Reduced AHP following BKCa1.1 LOF reported here is similar to that observed following inhibition of BK in cerebellar Purkinje cells that also increased burst firing(*21*). Based on these observations, we propose that BKCa1.1 channels terminate burst events through regulation of the action potential afterhyperpolarization in VTA dopamine neurons, and in their absence phasic dopamine is enhanced, which assigns a higher value to action and the outcome. This facilitates the acquisition of the conditioned response and establishes a greater differential response during extinction; thus, enhancing the reward prediction error and driving a larger extinction burst (*2, 3, 22, 23*).

We find that Kv4.3 channels regulate firing frequency, likely in response to descending GABAergic disinhibitory mechanisms (*24*), through the regulation of membrane repolarization(*9*). In the absence of this inhibitory influence, tonic signals such as ramping are elevated and enhance performance of a learned response, consistent with previous assertions on the function of ramping (*3–6*). In contrast to previous reports that ramping of dopamine release occurs independent of activity at the cell body (*5*), our data demonstrate that the intrinsic excitability of the cell, regulated by Kv4.3, contributes at least partially to this process in reinforced trials. During the first day of extinction training, Kv4.3 LOF mice had a lower extinction behavior burst than control or BKCa1.1 LOF mice. We also observed that Kv4.3 LOF mice had reduced dopamine neuron action potential firing of dopamine neurons and reduced dopamine release. This suggests that responding during the lever press period and during the reward omission period both contribute to the behavior observed during extinction. Interestingly, we did observe a disconnect between the dopamine neuron action potential firing and dopamine release in Kv4.3 LOF mice in the lever press period during extinction. Action potential firing was neither increased nor decreased; however, dopamine release dipped below baseline. This suggests that terminal modulation through inhibition of release may occurring during extinction and that this effect is unmasked in Kv4.3 LOF mice that do not display an increase in action potential firing which would partially counter this inhibition. The source of this inhibitory signal is not clear but may involve dynorphin release within the NAc that acts on kappa opioid receptors expressed on dopamine terminals (*31, 32*), which is known to modulate reinforcement behavior (*33*).

A limitation of the current study relating to the loss of function of Kv4.3 and BKCa1.1 is the potential impact on conductance associated with other ion channels. Indeed, as mentioned above, BK currents can influence Kv2 currents, and vice versa in SNc dopamine neurons (*9*). Because Kv4 channels are activated during rapid ramping of the depolarization they help to constrain the inward sodium current; thus, their loss would be predicted to result in a larger sodium conductance and higher frequency firing (*9*). In VTA dopamine neurons a small subthreshold sodium current has also been shown to contribute to pacemaker activity (*23*), reducing the resting potential to varying degrees following Kv4.3 or BKCa1.1 mutagenesis could disrupt the subthreshold sodium conductance and alter the excitability of VTA dopamine neurons or their pacemaker activity. These studies highlight the critical interactions between ion channels that work collectively to regulate action potential firing. Regardless of the mechanism(s) that underlies the impact of Kv4.3 and BKCa1.1 LOF on VTA dopamine neuron physiology, we find significant effects on the action potential waveform, action potential firing both *ex vivo* and *in vivo*, and changes in dopamine release dynamics that supports the importance of their cell autonomous function in these cells.

Although we focused on Kv4.3 and BKCa1.1 in dopamine neurons, these channels are expressed widely throughout the brain where they likely serve similar functions in patterning transmitter release. Further, there are numerous other ion channels that contribute to the action potential. The emergence of many of these ion channels as hotspots of missense mutation in neurodevelopmental disorders (*15*) reinforces the importance of understanding how these channels influence cellular and systems function.

## Materials and Methods

### Experimental Design

#### Mice

All procedures were approved and conducted in accordance with the guidelines of the University of Washington’s Institutional Animal Care and Use Committee. Mice were housed on a 12:12 light: dark cycle with *ad libitum* access to food and water, except when undergoing food restriction for Pavlovian and operant behavioral conditioning. Approximately equal numbers of male and female mice were used. Mice were group-housed (2-5 mice per cage), except during night-day locomotion assays. Either C57BL/6 (for *in situ* hybridization with RNAscope) or *Slc6a3*-IRES-Cre (DAT-IRES-Cre (Jackson Laboratory, Jax 006660), backcrossed to C57BL/6) mice were used.

#### Viruses

All adeno-associated viruses (AAV) were produced in-house with titers of 1-3X10^12^ particles mL^-1^, as described (*25*). Cre-inducible CRISPR viruses: AAV1-CMV-FLEX-SaCas9-U6-sgKcnd3, AAV1-CMV-FLEX-SaCas9-U6-sgKcnma1. Cre-inducible optogenetic viruses: AAV1-EF1α-FLEX-ChR2-eYFP and AAV1-EF1α-FLEX-Chrimson-TdTomato. Dopamine sensor virus: AAV1-CAG-dLight1.3B. Cre-inducible control virus: AAV1-EF1α-FLEX-eYFP. Cre-inducible FACS virus: AAV1-hSyn1-FLEX-KASH-GFP.

#### CRISPR/Cas9 design

sgRNAs were designed using methods previously described (*13, 26*). 21 bp sgRNAs for *Kcnd3* (Forward: GAGCGCTATCCCGACACCTTG; Reverse: CAAGGTGTCGGGATAGCGCTC) and *Kcnma1* (Forward: GTCTAGGCTGAGATGGTTCGC; Reverse: GCGAACCATCTCAGCCTAGAC) were generated (Invitrogen) and cloned into pAAV-CMV-FLEX-SaCas9-U6sgRNA for generation of AAVs in house.

#### Surgeries

DAT-IRES-Cre mice were 6-8 weeks of age at time of surgery, except for mice used for in vivo electrophysiology which were at least 10 weeks of age at time of surgery. Mice were anesthetized with isoflurane in a chamber and then placed and secured into the stereotax with isoflurane delivery through a nose cone. For all implants, an anchoring screw was driven into the skull away from the implant site. Mice received eYFP (control), CRISPR/eYFP (*in vitro* electrophysiology or behavioral groups), CRISPR/KASH-GFP (sequencing) or CRISPR/ChR2 (*in vivo* electrophysiology) bilateral injections into the VTA (in mm, relative to bregma: A/P: - 3.3; M/L ± 0.5; D/V: -4.9 - -4.4), total volume 0.5 µL into each side. A/P coordinates were adjusted for Bregma-Lambda distances using a correction factor of 4.21 mm. For D/V, the syringe was lowered to the deepest indicated coordinate and then raised at the start of the injection. For ex vivo two-photon dLight experiments, all mice received a bilateral injection ofdLight1.3b into the NAc (in mm, relative to bregma: A/P: +1.42; M/L ± 0.9; D/V: -4.5➔-4.1) followed by bilateral VTA injections of CRISPR and Chrimson (0.5 µL each side; control mice received Chrimson viruses diluted to the same ratio). For in vivo fiber photometry of dLight, mice received control or CRISPR viruses bilaterally into the VTA and then were injected unilaterally with dLight1.3b into the NAc (in mm, relative to bregma: A/P: -+1.43; M/L - 0.9; D/V: -4.5➔-4.1), total volume 0.5 µL; a photometry fiber purchased from Doric (400 µm optical fiber, 1.25 mm metal ferrule) was lowered to D/V -4.1 mm. Fibers were secured to the skull with Metabond (Parkell) and dental cement (Lang Dental). For *in vivo* electrophysiology, microdrives were constructed in-house with a Neuralynx EIB (EIB-16) mounted on a 3D printed L-frame (Applied Rapid Technologies). Each microdrive contained in house constructed optetrodes of an in house constructed optic fiber (200 µm fiber bore and 1.25 mm ceramic ferrule, Precision Fiber Products) and an in house constructed tetrode bundle of tungsten wire (each tetrode was 4 wires 25 mm diameter spun together and heat-sealed, California Fine Wire). After virus injection for CRISPR/ChR2 mixture, a window was created in the skull and the microdrive was lowered to above the VTA (−3.8 mm). Microdrives were secured to the skull with Metabond and dental cement.

#### *In situ* hybridization using RNAscope

Wildtype C57BL/6 male and female mice (8-9 weeks old) were used to map ion channel expression in the VTA using RNAscope (ACDBio) (*27*). Brains were flash frozen in 2-methylbutane, and stored at -80LJ. VTA sections were collected and processed for RNAscope (20 um, coronal). Representative sections that spanned the VTA were selected for hybridization (A/P in mm: rostral: -3.1; intermediate: -3.5; caudal: -3.8; according to the Paxinos atlas). Sections were prepared for hybridization per manufacturer’s (Advanced Cell Diagnostics, Inc) instructions using probes for *Th* (Mm-*Th*), *Kcnd3* (Mm-*Kcnd3*-C3), and *Kcnma1* (Mm-*Kcnma1*-C2). Slides were coverslipped with Fluoromount with DAPI (Souther Biotech) and imaged using a confocal fluorescent microscope (University of Washington Keck Center (NIH grant S10 OD016240), Leica SP8X confocal). Analysis was performed Cell Profiler 4.1.3 (https://github.com/CellProfiler/CellProfiler) using a modified image analysis pipeline for RNAscope (*28*). Images for each RNAscope experiment were collected using the same acquisition settings for each slide.

#### Immunohistochemistry for target validation

Following completion of experiments, all mice were checked for viral targeting or implant placement. Mice were deeply anesthetized with Buthenesia and transcardially perfused with cold 1X PBS followed by cold 4% paraformaldehyde (PFA, Sigma-Aldrich). Implants were removed from mice used in *in vivo* electrophysiology or fiber photometry experiments. Brains were post fixed for 24 hours in PFA at 4LJ, then moved to 30% sucrose in PBS until cryosectioning occured. All cryosections were stored in 1XPBS with 0.3% Sodium Azide (Sigma-Aldrich) until immunohistochemistry. For colocalization validation and behavioral experiments, the VTA was coronally sectioned at 30 µm. For *in vivo* electrophysiology experiments, the VTA was coronally sectioned at 60 µm. For *in vivo* dLight fiber photometry experiments, coronal sections of the NAc (60 µm) and the VTA (30 µm) were obtained. For *ex vivo* dLight slice experiments, the part of the brain containing the VTA was blocked at the same time as obtaining NAc slices for slice imaging. The block was post fixed overnight in 4%PFA at 4LJ. They were then transferred to 30% sucrose in PBS for at least 24 hours and then VTA slices were cryosectioned at 30 µm. NAc slices (250 µm) used in dLight slice two-photon imaging were immediately transferred to 4%PFA after recording and were fixed overnight at 4LJ. Slices were then transferred to 1X PBS until day of immunohistochemistry. Free floating immunohistochemistry experiments were performed using the following anitbodies as needed: rabbit anti-HA (Sigma), mouse anti-TH (Millipore), mouse anti-dsred (Takara), rabbit anti-GFP (Proteintech), and chicken anti-GFP (Aves) to target CRISPR, TH, Chrimson-TdTomato, Chr2-eYFP, or dLight1.3-GFP virues followed by secondary antibody application (JacksonImmuno). Sections were placed on slides, coverslipped with Fluoromount with DAPI, and kept in the dark at 4LJ until the day of imaging. Images were collected at the UW Keck Center or on a Keyence Fluorescence Microscope (Keyence). Quantification of co-labeled cells for immunohistochemistry was performed using ImageJ 1.53 Cell Counter/Multi-point tool.

#### Fluorescence-activated cell sorting and targeted deep sequencing

Procedures were performed as previously described (*13, 26*). AAV1-hSyn1-FLEX-EGFP-KASH and CRISPR viruses (AAV1-CMV-FLEX-SaCas9-U6-sgRNA) targeting either *Kcnd3* or *Kcnma1* were stereotaxically co-injected into the VTA of DAT-IRES-Cre mice. Four weeks after surgery, the VTA of each mouse was extracted, flash frozen in liquid nitrogen, and stored at -80°C. On day of FACS, VTA tissue from 5 mice per experimental group were combined into a single sample and homogenized in buffer containing (in mM): 320 Sucrose (sterile filtered), 5 CaCl (sterile filtered), 3 Mg(Ac)2 (sterile filtered), 10 Tris pH 7.8 (sterile filtered), 0.1 EDTA pH 8 (sterile filtered), 0.1% NP40, 0.1 Protease Inhibitor Cocktail (PIC, Sigma), 1 β-mercaptoethanol. Nuclei were isolated by centrifugation in 29% iso-osmolar Optiprep solution. GFP+ and GFP- nuclei were sorted using a BD AriaFACS III (University of Washington Pathology Flow Cytometry Core) into a PCR tube strip containing 3 µL of REPLI-g Advanced Single Cell Storage buffer (Qiagen). Whole genome amplification (WGA) was performed directly following FACS using the REPLI-g Advanced DNA Single Cell Kit (Qiagen) according to manufacturer’s instructions. For generation of amplicons, samples were amplified (PCR 1) with Phusion High Fidelity Polymerase (Thermo Fisher) followed by reamplification (PCR 2) with a second set of primers. The amplicons were gel extracted using the MinElute gel extraction kit (QIAGEN) and sent to Genewiz for Sanger and amplicon sequencing (Amplicon-EZ). The following primer sets were used to generate <300bp amplicons: PCR 1: *Kcnd3* forward: CATTGGATGGATGCCAGT; *Kcnd3* reverse: GAGGATGCCATAGAAGGC; *Kcnma1* forward: GCAACATGGCTGTTGATG; *Kcnma1* reverse: ACCTCCATGGTCACCGGT. PCR 2: *Kcnd3* forward: TAGCTCCAGCGGACAAGA; *Kcnd3* reverse: CAGAGATGCATTCATAGC; *Kcnma1* forward: CCATTGCTAGCTATGGCA; *Kcnma1* reverse: ATGAGCGCATCCATCTTG.

Quantitative PCR (qPCR): qPCR was utilized to determine mRNA expression of relevant potassium channel subunits in the VTA of control, Kv4.3 LOF and BKCa1.1 LOF mice. Four weeks after CRISPR injection, tissue punches containing the VTA (bilateral) were collected and immediately frozen on dry ice, prior to storage at -80°C. RNA isolations were conducted from tissue samples using Qiagen RNeasy Micro Kit (Qiagen, Catalog no. 74004) according to manufacturer’s instructions. The RNA quality and concentration was assessed for each sample spectrophotometrically (A260/A280 & gt;1.9) by a NanoDrop instrument. Then 100 ng of RNA was reverse transcribed by PCR using SuperScript IV VILO (Invitrogen, Catalog no. 11755050). qPCR was conducted on the cDNA using a Taqman Gene Expression assay consisting of the Taqman Gene Expression Master Mix, and the appropriate Taqman probes in FAM channel (Master Max: Thermo Fisher Scientific, Applied Biosystems, catalog no. 4369016, Taqman probes: *GAPDH* (Assay ID: Mm99999915_g1; Cat. number: 4448489), *En1* (Assay ID: Mm00438709_m1_g1; Cat. number: 4448892), *Th* (Assay ID: Mm00447557_m1_g1; Cat. number: 4331182), *Kcnn3* (Assay ID: Mm00446516_m1; Cat. number: 4448892) and *Kcnd2* (Assay ID: Mm01161732_m1; Cat. number: 4448892). All qPCRs were performed in triplicates and run on an Agilent Technologies MX3005P qPCR machine at 50°C for 2 mins, 95°C for 20 sec, then 95°C for 3 sec and 60°C for 30 sec for 40 cycles. The threshold cycle (Ct value) of each gene was then normalized to GAPDH. ΔCT values were calculated by subtracting the geometric mean of CT values for *Gapdh*, *En1*, and *Th* from the CT values of the gene of interest (*Kcnd2* or *Kcnn3*). Control ΔCT values for the gene of interest were subtracted from the ΔCT values of the experimental group to generate ΔΔCTs. Fold change was calculated as 2^-(ΔΔCT).

#### Slice preparation for *ex vivo* physiology

Mice injected with CRISPR/eYFP or CRISPR/Chrimson/dLight1.3b were allowed 4-5 weeks recovery after surgery to allow for viral expression, mutagenesis, and residual protein turnover. All solutions were continuously bubbled with O_2_/CO_2_. Coronal brain slices containing the VTA (200 µm for VTA physiology) or NAc (250 µm for dLight slice two-photon imaging) were prepared in a slush NMDG cutting solution (in mM: 92 NMDG, 2.5 KCl, 1.25 NaH_2_PO_4_, 30 NaHCO_3_, 20 HEPES, 25 glucose, 2 thiourea, 5 Na-ascorbate, 3 Na-pyruvate, 0.5 CaCl_2_, 10 MgSO_4_, pH 7.3–7.4 (*29*). Slices recovered for ∼12 min in the same solution in 32°C water bath. Slices were then transferred to a room temperature HEPES-aCSF solution (in mM: 92 NaCl, 2.5 KCl, 1.25 NaH_2_PO_4_, 30 NaHCO_3_, 20 HEPES, 25 glucose, 2 thiourea, 5 Na-ascorbate, 3 Na-pyruvate, 2 CaCl_2_, 2 MgSO_4_). Slices recovered for an additional 60 min.

#### Slice electrophysiology

Whole-cell patch-clamp recordings were made using an Axopatch 700B amplifier (Molecular Devices) with sampling at 10KHz and filtering at 1 kHz. eYFP+ cells were visually identified via fluorescence for patching with electrodes at 3–5 MΩ. Series resistance was monitored during all recordings with changes in resistance +10% qualifying the cell for exclusion. Recordings were made in aCSF (in mM: 126 NaCl, 2.5 KCl, 1.2 NaH_2_PO_4_, 1.2 MgCl_2_ 11 D-glucose, 18 NaHCO_3_, 2.4 CaCl_2_) at 32°C continually perfused over slices at a rate of ∼2 ml/min. VTA dopamine neurons were identified by fluorescence. Electrodes were filled with an internal solution containing (in mM): 130 potassium gluconate, 10 HEPES, 5 NaCl, 1 EGTA), 5 Mg2+/ATP, 0.5 Na+-GTP, pH 7.3, 280 mOsmol.

#### Spontaneous activity, excitability, and action potential shape measurements

Spontaneous activity was recording in I=0 mode for 1 minute and I/V curves (0-80 pA, 10 pA steps, 1 s) for excitability were immediately measured in I-Clamp mode (done both at cell’s resting membrane potential and when holding the cell at -60 mV). All data were analyzed offline using Clampfit (Molecular Devices). Frequency of spontaneous activity was calculated as number of events/s and CV-ISI was measured using the standard deviation of the interspike interval (ISI) from spontaneous activity. Excitability was measured as the total number of events during each current step. Action potentials of spontaneous activity were detected using the Event Detection function in Clampfit (restricted between 50 ms before action potential peak and 150 ms after action potential peak). The first action potentials of activity elicited from a 20 pA current injection while the cell was being held at -60mV were also detected using the Event Detection function in Clampfit with identical parameters. Event statistics from the Event Detection function were extracted, including action potential peak, half-width, decay tau, and afterhyperpolarization (the anti-peak Event Statistics characteristic corrected for resting membrane potential measure 50 msec before spike peak). For each cell, detected action potentials were averaged to visualize action potential waveforms and phase plane plots. Phase plane plots were calculated as the dV/dt and plotted as the dV/dt over voltage. Action potential threshold was defined as the point at which dV/dt of the AP waveform reaches 4% of its maximum (*23*). Resting membrane potential was recorded as the membrane potential at the beginning of the action potential waveform from the Event Detection window (50 msec before peak). For action potential shape analysis of control VTA dopamine neurons following pharmacological inhibition of A-Type or BK potassium currents, AmmTx3 (10 nM, Tocris) and Iberiotoxin, (150 nM, Cayman Chemical) were perfused onto the slice in oxygenated aCSF and spontaneous activity was recorded. Shape analyses were performed as described above.

#### Potassium current measurements

For A-Type and BK potassium channel current, *I*_A_ and *I*_BK_ respectively, recordings in V-Clamp mode, aCSF was supplemented with 500 nM TTX to eliminate action potential activity and cells were held at -60 mV. *I*_A_ was activated using voltage steps from -108mV to -28 mV to -48mV then back to holding potential every 5 s, 5 times. AmmTx3 (10 nM, Tocris) was used to pharmacologically block Kv4 channels. *I*_BK_ was measured using subtraction before and after pharmacological inhibition of BK channels (iberiotoxin, 150 nM (Cayman Chemical)) during depolarizing voltage steps that activated *I*_K_ from -63mV to +2mV to -48 mV to -63 mV every 5 s 5 times. Data were analyzed offline using Clampfit. For A-type potassium currents, differences between the peaks at the -28mV and the -48 mV step were determined and calculated. For BK currents, peaks and area under the curves of *I*_K_ before and after iberiotoxin was determined and then the subtracted current pre- and post pharmacological inhibition was extracted. Maximal peaks and AUC were then calculated for the subtracted current to derive *I*_BK_.

#### *In vivo* single-unit electrophysiology

Baseline recordings began 4 weeks after surgery in a recording cage with corncob bedding. Mice were allowed to freely roam in the cage during recording, no nestlet was provided. For *in vivo* electrophysiology recordings during behavior (behaviors described below), mice were recorded in operant chambers (MedAssociates) that were interfaced with the Data Acquisition software (Digital Lynx 4SX, Neuralynx). For all experiments, the ceramic ferrule of the optic fiber in the microdrive was attached to a patch cord that was connected to a 473 nm blue laser (Laserglow technologies). A preamplifier was connected to the microdrive’s EIB. Recordings were acquired through Cheetah Data Acquisition and optical stimulations were synced through Cheetah with a function generator (Agilent). Data were continuously sampled, amplified, and filtered between 100 and 9000 Hz, and digitized at 32 kHz. If spike exceeded a predetermined threshold, discretely sampled data were recorded and filtered between 100 and 6000 Hz for 10-20 minutes. At the end of the recording, 10 blue light pulses were delivered 20 times through the optic fiber of the microdrive to activate neurons that expressed ChR2 (5-ms long at 20 Hz, putative dopamine neurons). The light intensity (5-15 mW/mm2) was adjusted so that light-evoked spike waveforms were similar to spontaneous ones. All optetrodes were lowered in 80 um increments at the end of the day to identify a new cell field the next day. Data were analyzed offline using Plexon Offline Sorter, SpikeSort3D, Neuroexplorer, and Matlab, as previously described (*30*). Briefly, units were identified using cluster analysis in the Offline Sorter or SpikeSort3D (KlustaKwik) and saved. Frequency, burst analysis, and blue-light sensitive (ChR2 expressing putative dopamine neurons) identification were performed in Neuroexplorer and Matlab. Light sensitive units were identified as those that had a spike latency of ≤ 5 s and 0.6 probability during blue light activation of ChR2. The onset of a burst was identified as two consecutive spikes with an interspike interval of <80 ms and its termination was defined as an interspike interval of >160 ms (*30*). For analysis of in vivo firing of optically-identified dopamine neurons, ± 10 s perievent windows for relevant behavior events (lever press (LP), reward delivery (Rew), and head entry/lever extension (HE/CS)) were extracted and a z-scores calculated; AUCs were then determined for these events: LP (3 s post), Rew (5 s post), HE/CS (5 s post).

#### Overnight locomotor activity

4 weeks after surgery, baseline locomotion was measured using locomotion chambers (Columbus instruments) that use infrared beam breaks to calculate ambulatory activity. Mice were singly housed and provided with *ad libitum* access to food and water. Locomotion was monitored for 60 consecutive hours (Friday-Monday: 3 nights, 2 days). Behavioral tracking failed for one BKCa1.1LOF mouse, however this mouse was still included in operant studies.

#### Pavlovian conditioning

At least 4 weeks after surgery, mice were food restricted to 85% of their body weight and were maintained at this bodyweight for all conditioning days. Conditioning sessions took place in mouse conditioning boxes outfitted with a houselight, ultrasensitive levers, a food hopper with a head entry detector, nose poke holes, and barred floors (MedAssociates). Pavlovian training sessions consisted of 25 trials in which 10 s lever extension (an audiovisual CS). After 10 s lever presentation, levers were retracted and a sucrose pellet reward (Rew, 20 mg sucrose pellet, Bioserv) was delivered in the food hopper. Each trial was separated by a random inter-trial interval averaging 70 s and animals underwent 7 days of training. The houselight remained on during the entire session. Conditioned approach scores were calculated as the rate of head entries during the CS - rate of head entries during the ITI. Mice used for *in vivo* electrophysiology or dLight experiments first underwent 3 days of 20 min acclimation sessions to patch cord tethering in the conditioning boxes followed by 10 days of Pavlovian conditioning while tethered.

#### Operant conditioning

After Pavlovian conditioning, mice began an FR1 schedule of reinforced operant conditioning. Each session lasted for 60 min. Here, levers were extended and remained extended until a LP. Upon a lever press, levers were retracted, and a sucrose pellet was immediately delivered into the food hopper (Rew). For *in vivo* electrophysiology or dLight experiments during FR1 reinforced training, a temporal delay of 3 s between LP and Rew was included to dissociate dynamics of LP and Rew. The levers did not extend again until the mouse entered the food hopper to retrieve the sucrose pellet (HE/CS). The house light remained on during the entire session. Reinforced FR1 sessions lasted for 3 days (5 days for dLight mice). A subset of mice then underwent a progressive ratio test where the number of lever presses necessary for sucrose pellet delivery increases non-arithmetically (i.e., 1, 2, 4, 6, 9, 13…) over the course of the session. The progressive ratio session automatically ended after 3 consecutive min of no lever presses or after 3 hours. Another subset of mice that had undergone FR1 reinforced training underwent FR1 extinction for 60 min each session for five days. Here, levers extend and retract similarly to the FR1 reinforced paradigm, yet a sucrose pellet reward is omitted. Mice underwent extinction for 5 days. *In vivo* electrophysiology and dLight mice also underwent 5 days of extinction training following completion of FR1 reinforced training.

#### dLight recordings using fiber photometry during operant conditioning

Fiber photometry recordings lasted the entirety of the behavioral conditioning sessions (first and last days of reinforced FR1 and extinction FR1). dLight1.3 was used to measure dopamine release in the NAc during these behaviors of control or CRISPR mice. Prior to every session, the implanted optic fiber was attached to a previously photobleached patchcord (Doric Lenses) using a ceramic sleeve (Doric Lenses). Recordings were acquired using an RZ5 BioAmp Processor from Tucker Davis Technologies (TDT). A 465 nm LED (531-Hz, sinusoidal, Doric Lenses) was used to excite dLight1.3b and a 405 nm LED was used as an isosbestic control (211-Hz, sinusoidal, Doric Lenses) using an LED driver (Doric Lenses) controlled via the RZ5. Prior to each session, patch cords had been photobleached for at least one hour. Laser intensities were measured at the tip of the optic fiber to reach levels of 30-45 uW before being bandpass filtered (525 ± 25 nm, Doric, FMC4), transduced by a femtowatt silicon photoreceiver (Doric Lenses), and recorded by a real-time processor (TDT, RZ5). The 531-Hz and 211-Hz signals were extracted in real-time by the TDT program Synapse at a sampling rate of 1017.25 Hz. MedAssociate signals of behavioral events such as LP, Rew, and HE/CS were synced to the recordings for further analysis offline.

#### Photometry analysis of perievents

A custom software GUI was generated to extract, downsample and z-score the photometry signals surrounding behavioral events for each mouse (20 s window: ± 10 s behavioral event). Data were first downsampled to 100 Hz and DF/F was calculated using the custom software GUI. Next, Z-score was calculated relative to the mean and standard deviation of the first 4 s of the 20 s time window (baseline perievent time). Mouse data were further processed in custom MatLab script. Z-scores for each mouse were then downsampled to 50 Hz and then averaged across all behavioral. Cumulative summation of dopamine ramp preceded LP for the 10 s preceding the behavioral event. Data were exported to GraphPad Prism for statistical analysis, area under the curve (AUC) analysis, and trace export. For reinforced FR1 training and omission extinction training, AUC time bins were calculated for a set time following the event: LP (3 s post), Rew (5 s post), HE/CS (5 s post). For extinction, the AUC for HE/CS was divided into two phases following the event: period A (0-2 s post) and period B (2-5 s post).

#### Two(2)-photon slice imaging of dLight

NAc slices were transferred to a heated recording in aCSF (in mM: 126 NaCl, 2.5 KCl, 1.2 NaH_2_PO_4_, 1.2 MgCl_2_ 11 D-glucose, 18 NaHCO_3_, 2.4 CaCl_2_) at 32°C continually perfused over slices at a rate of ∼2 ml/min on an Olympus Fluoview FVMPE-RS 2-photon microscope.. The NAc was identified with epifluorescence and expression of dLight GFP and VTA dopamine terminals expressing Chrimson-TdTomato was quickly confirmed before recording. We used a resonant scanner (30 Hz frame rate acquisition) and performed an online averaging of 6 times to get an effective frame rate of 5 Hz. A GaAsP-PMT with adjustable voltage, gain and offset was used, along with a green filter cube. We used a 25x immersion objective (Olympus, XLPLN25XWMP, 1.05 NA, 2 mm WD) and imaged with a **920** nm laser (SpectraPhysics, ∼100 fs pulse width) with automated wavelength-specific alignment. Chrimson fibers in the NAc were stimulated at increasing frequencies for 1 sec following a 10 sec baseline using full field 615 nm LED stimulation through the objective (CoolLED, PE-100). Data were analyzed in FiJi ImageJ using the Time Series Analyzer plug in. Z-score of fluorescence was calculated from a 10 second baseline preceding the Chrimson stimulation. AUC calculations were performed in GraphPad Prism.

#### Statistics

Data were analyzed for statistical significance using GraphPad Prism. All statistical tests were two-sided and corrected for multiple comparisons where appropriate. Detailed statistical analyses can be found in the figure legends.

## Acknowledgments

We would like to thank members of the Zweifel, Palmiter, and Chavkin laboratories at the University of Washington for their helpful discussions. We would also like to thank the staff of the University of Washington’s Molecular Genetics Research Core, Imaging and Neural Circuits Core, Comparative Medicine Animal Facilities, the University of Washington’s Keck Imaging Center, and the University of Washington’s Pathology Flow Cytometry Core.

## Funding

This study was supported by:

K99-DA054265 (B.J.)

T32-DA727825 (B.J.)

F32 MH127801 (M-S.K)

F31 DA053724 (J.E.E)

F31-MH116549 (A.C.H.)

T32-GM007270 (A.C.H.)

F31 MH126489 (M.C.)

T32-GM007270 (M.C.)

R03TR003307 (M.E.S.)

R01-MH104450 (L.S.Z.)

R01-DA044315 (L.S.Z.)

University of Washington’s Addictions, Drugs and Alcohol Institute:

ADAI-1019-17 (B.J.)

National Research Foundation of Korea:

MEST 2020R1A2C2102134 (Y.S.J)

Core grants:

S10OD016240 (University of Washington Keck Center)

P30-DA048736 (University of Washington Center of Excellence in Opioid Addiction Research) B.J. was funded by the postdoctoral Enrichment Program Award from the Burroughs Wellcome Fund.

## Author contributions

Conceptualization: BJ, LSZ

Methodology: BJ, MES, SN-E, LSZ

Investigation: BJ, M-SK, YSJ, JEE, JXY, MC, ACH, MAQ, MAB, MJ, DB AM, NLG

Visualization: BJ, ACH, LSZ

Supervision: BJ, LSZ

Writing—original draft: BJ, LSZ

Writing—review & editing: all authors reviewed and approved this manuscript. BJ and LSZ edited the manuscript following co-author review.

## Data and materials availability

Data and code can be made available upon request.

## Supplementary Materials

Supplementary figures.

