## Supplemental File for "Temporal scaling of dopamine neuron firing and dopamine release by distinct ion channels shape behavior"

### Supplemental Materials
### Supplemental figures and supplementary figure legends:

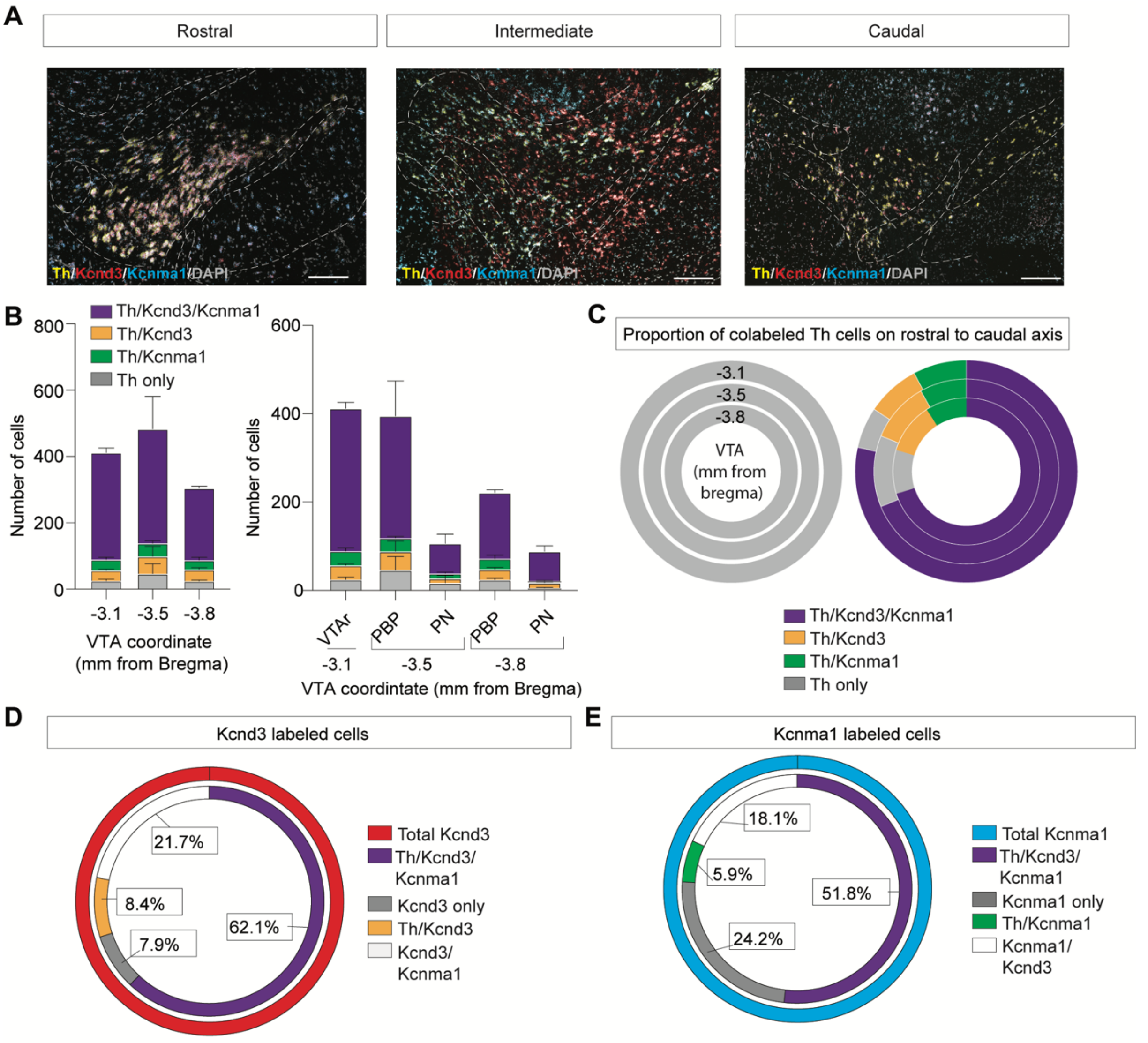


| Total Kcnd3 | 4275 |
|---|---|
| Kcnd3/Th/Kcnma1 | 2653 |
| Kcnd3 only | 337 |
| Kcnd3/Th | 358 |
| Kcnd3/Kcnma1 | 927 |

| Total Kcnma1 | 5121 |
|---|---|
| Kcnma1/Th/Kcnd3 | 2653 |
| Kcnma1 only | 1238 |
| Kcnma1/Th | 303 |
| Kcnma1/Kcnd3 | 927 |

**Fig S1: In situ hybridization (RNAscope) of Th, Kcnd3, and Kcnma1 in the adult mouse VTA.**

(**A**) Representative in situ images of coronal VTA sections probed for Th, Kcnd3, and Kcnma1. (**B**) Quantification of cell overlap in the VTA sections and subregions along the rostral-caudal extent of the VTA. VTAr: rostral VTA, PBP: parabrachial portion, PN: paranigral. (N=3 mice; Data presented as mean + SEM). (**C**) Proportion of total Th labeled cells and cell overlap with Kcnd3 or Kcnma1. (**D**) Proportion of total Kcnd3 labeled cells in the mouse VTA (top) and total cell counts (bottom table). (**D**) Proportion of total Kcnma1 labeled cells in the mouse VTA (top) and total cell counts (bottom table).

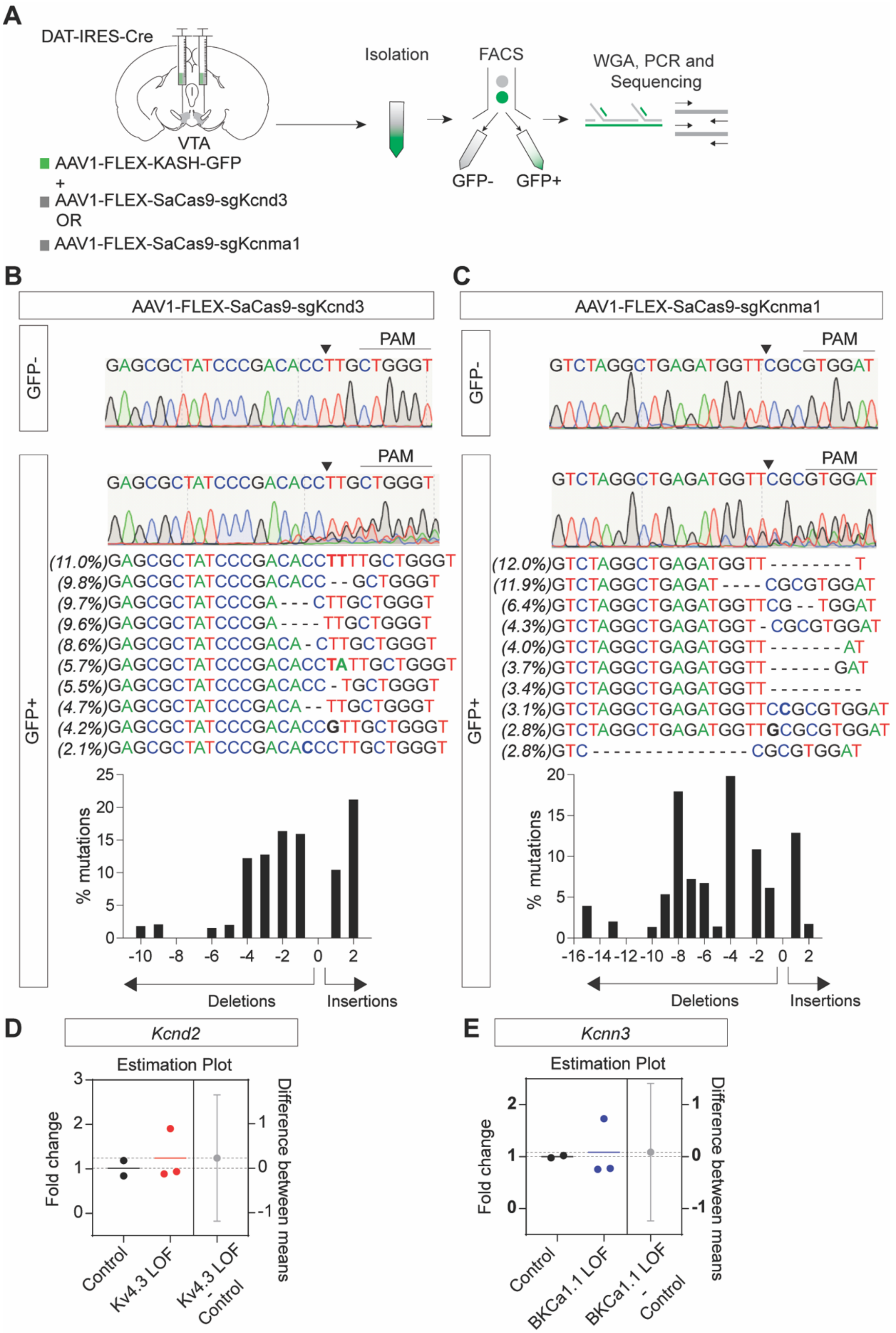


**Fig S2**: **Single vector Cre-inducible AAV CRISPR/Cas9 targeting of VTA dopamine neurons induces specific mutagenesis.**

(**A**) Experimental strategy to isolate KASH-GFP+ (putative dopamine) and GFP- (putative non-dopamine cells) in the mouse VTA of DAT-IRES-Cre mice for Sanger sequencing. (**B**) Quantification of mutagenesis in Kcnd3 gene in mouse VTA GFP- and GFP+ cells in AAV1-FLEX-SaCas9-sgKcnd3 injected mice (N=5 mice). (**C**)

Quantification of mutagenesis in Kcnma1 mouse VTA G(FP- and GFP+ cells in AAV1-FLEX-SaCas9-sgKcnd3 injected mice (N=5 mice). qPCR mRNA quantification of (**D**) *Kcnd2* in control and Kv4.3 LOF (Unpaired t-test, $t_{(3)=}$0.5137, P=0.6429; $N_{control}$=2 and $N_{Kv4.3LOF}$=3 mice) or (**E**) *Kcnn3* in control and BKCa1.1 LOF mice (Unpaired t-test, $t_{(3)=}$0.2085, P=0.8482; $N_{control}$=2 and $N_{Kv4.3LOF}$=3 mice). Data represented as mean $\pm$ SEM.

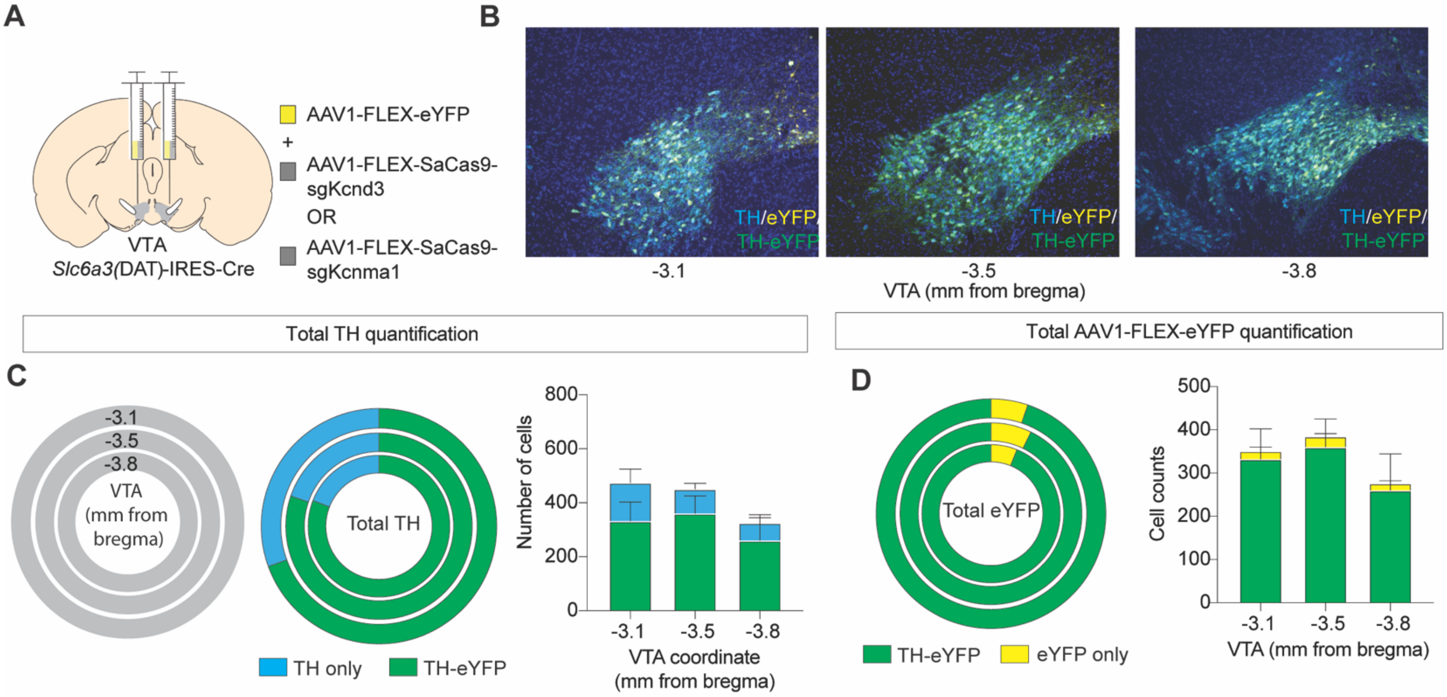


**Fig S3**: **Immunohistochemical analysis of viral transduction of dopamine neurons of the VTA.**

(**A**) Schematic of viral injection strategy for slice electrophysiology and behavioral cohorts. (**B**) Immunohistochemistry for eYFP and TH and 3 anatomical locations along the rostral-to-caudal axis. (**C**) Quantification of TH-positive neurons containing eYFP expression (N=3 mice). (**D**) Quantification of eYFP-positive neurons expressing TH (N=3 mice). Data in bar graphs presented as mean $\pm$ SEM.

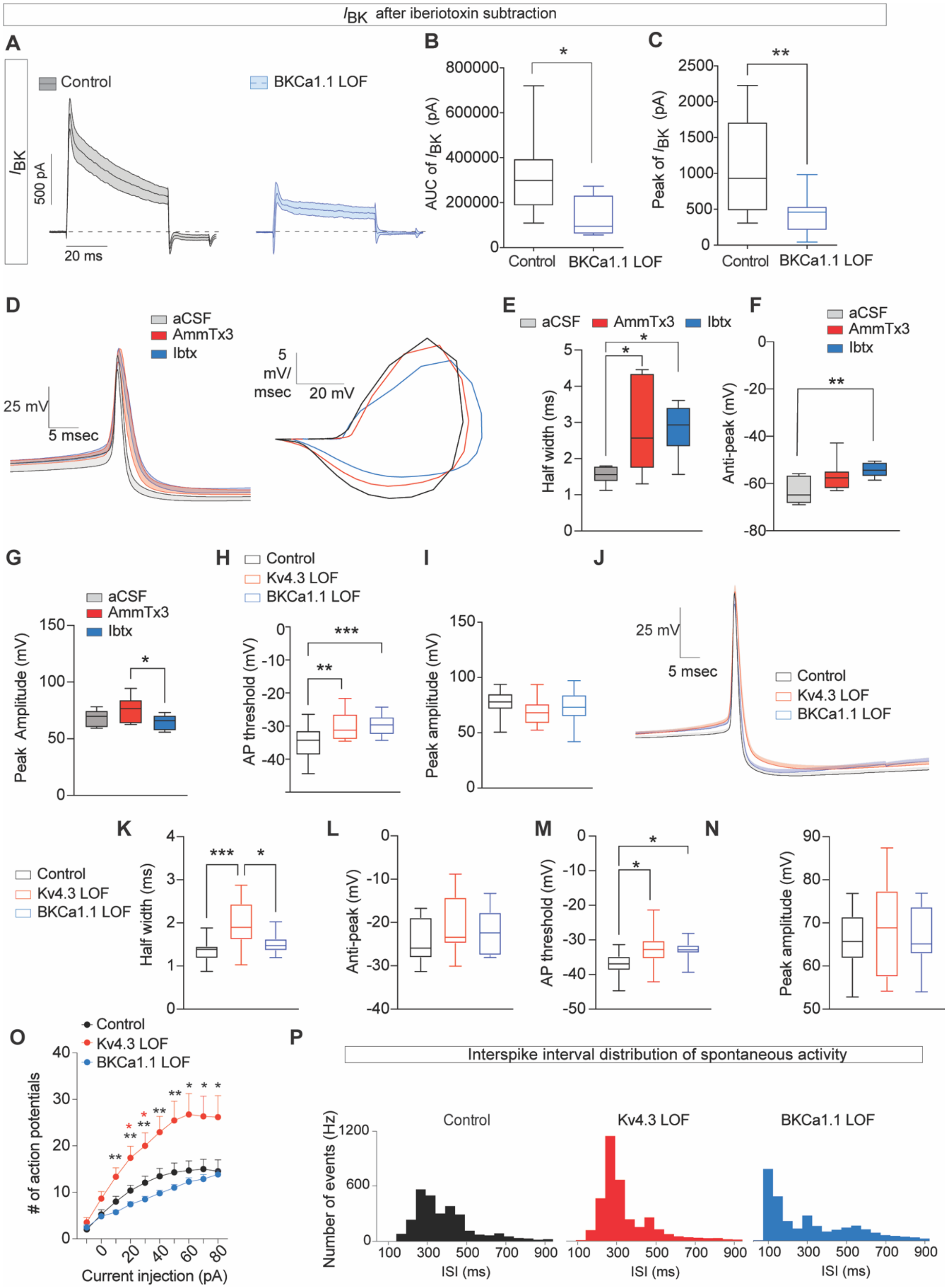


**Fig S4: Electrophysiological properties of Kv4.3 and BKCa1.1 LOF and pharmacological blockade of ion channels.**

(**A**) $I_{BK}$ after subtraction iberiotoxin-insensitive current. (**B**) sgKcnma1 significantly reduces the area under the curve (AUC) of $I_{BK}$ in BKCa1.1 LOF mice (Unpaired t-test, $t_{19}$=2.234, *P<0.05). (**C**) BKCa1.1 have significantly reduced subtracted peak of $I_{BK}$ (Unpaired t-test, $t_{19}$=3.738, **P<0.01)). (**B-C**, $N_{control}$=11 cells,

$N_{sgKcnma1}$=10 cells). (**D**) Action potential waveform (left) and phase plane plot (right) of spontaneously firing action potentials in aCSF, AmmTX3 (Kv4 channel blocker), and Iberiotoxin (Ibtx), BK channel blocker). (**E**) Action potential half width following pharmacological blockade (One-way ANOVA, $F_{(2,22)}$=5.266, P<0.05, Tukey's multiple comparison's test, *p<0.05). (**F**) Action potential after hyperpolarization (anti-peak) following pharmacological blockade (One-way ANOVA, $F_{(2,22)}$=6.040, P<0.01, Tukey's multiple comparison's test, **p<0.01). (**G**) Action potential peak amplitude following pharmacological blockade (One-way ANOVA, $F_{(2,22)}$=3.433, P=0.0504, Tukey's multiple comparison's test, *p<0.05). (**D-G**, $N_{control}$=7 cells, $N_{Kv4.3LOF}$=10 cells, $N_{BKCa1.1LOF}$=8 cells) (**H**) Action potential threshold of spontaneously active action potentials from control and CRISPR LOF cells (One-way ANOVA, $F_{(2,53)}$=11.09, P<0.0001, Tukey's multiple comparison's test, **p<0.01, ***p<0.01). (**I**) No differences in maximum peak amplitude from control and CRISPR LOF cells (One-way ANOVA, $F_{(2,53)}$=0.04907, P=0.9522). (**H-I**, $N_{control}$=21 cells, $N_{Kv4.3LOF}$=16 cells, $N_{BKCa1.1LOF}$=19 cells) of action potential shape from spontaneous activity. (**J**) Average action potential waveform of the first generated action potential from a 20 pA step. (**K**) AcEtion potential half width from first generated action potential from a 20 pA step (One-way ANOVA, $F_{(2,33)}$=9.263, P<0.001, Tukey's multiple comparison's test *p<0.05, **p<0.01). (**L**) Anti-peak (after hyperpolarization) from first generated action potential from a 20 pA step (One-way ANOVA, $F_{(2,33)}$=1.498, P=0.2383). (**M**) Action potential threshold from first generated action potential from a 20 pA step One-way ANOVA, $F_{(2,33)}$=5.289, P=0.0.104, P<0.05). (**N**) Peak amplitude from first generated action potential from a 20 pA step (One-way ANOVA, $F_{(2,33)}$=0.2903, P=0.7500). (**J-N**, $N_{control}$=13 cells, $N_{Kv4.3LOF}$= 12 cells, $N_{BKCa1.1LOF}$= 11 cells). (**O**) Input-output excitability curve from cells at resting membrane potential (Two-way RM ANOVA, Interaction effect $F_{(18,369)}$=4.027, P<0.0001; Group effect $F_{(2,41)}$=7.080, P<0.001; Current injection effect $F_{(1.238, 50.475)}$=53.90, P<0.0001; red asterisk=control vs Kv4.3 LOF, black asterisk= Kv4.3 LOF vs BKCa1.1 LOF, *p<0.05, **p<0.01). (**P**) Interspike interval mean time bins in spontaneous activity as a measurement of spike regularity. Box and whisker data are represented as min-to-max; action potential traces are presented as mean + SEM.

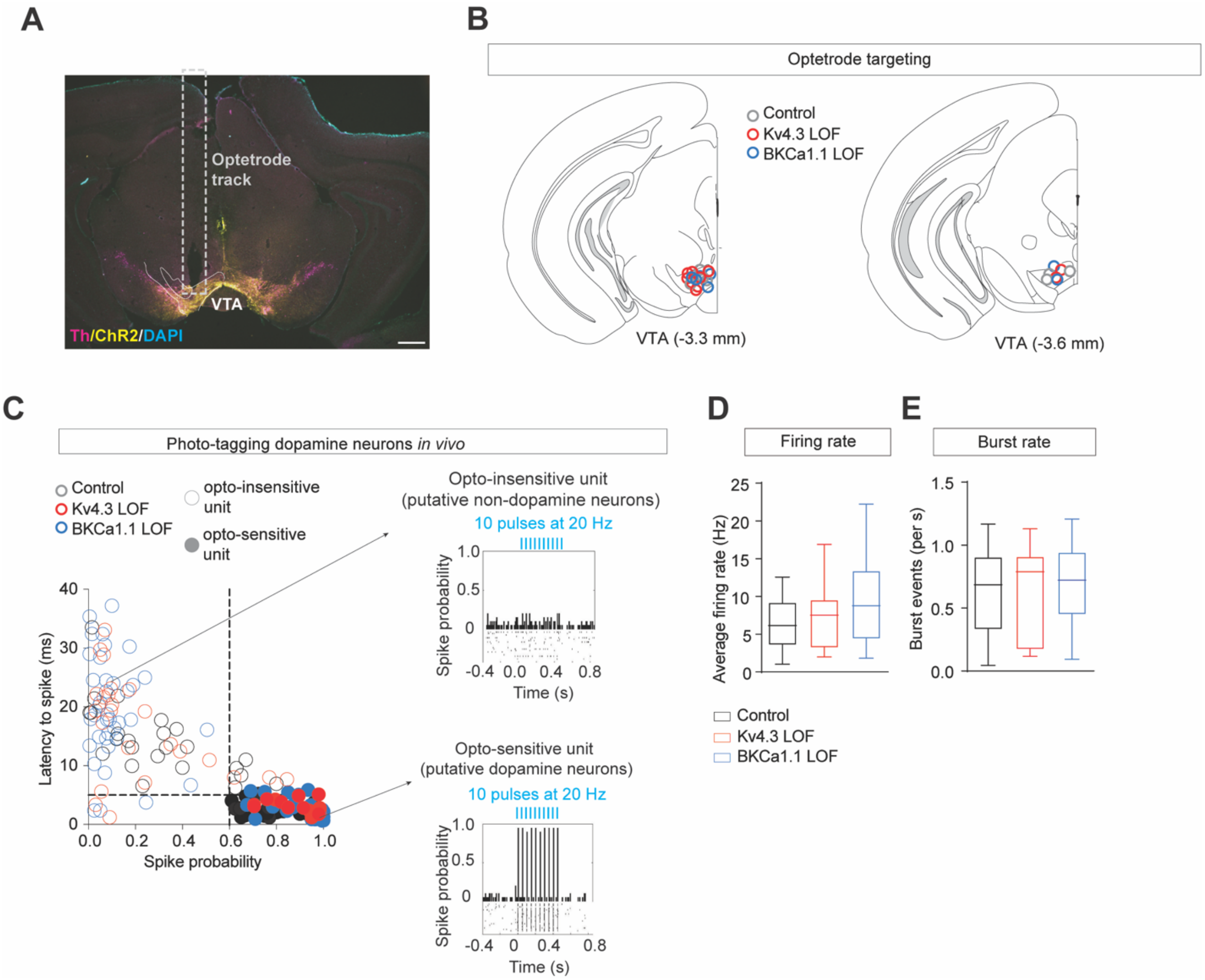


**Fig S5. Optetrode targeting and optogenetic identification of VTA dopamine neurons.**

(**A**). Representative image of optetrode track in adult mouse VTA with labeling of Th and virally expressed ChR2. Scale bar= 500 µm. (**B**) Targeting map for all recorded mice. (**C**) Latency vs spike probability plot of recorded units following optical excitation of ChR2 (left); units with spike latencies $\leq$5ms and spike probability >0.6 were considered opto-sensitive and putative VTA dopamine neurons. Opto-insensitive unit histogram during ChR2 excitation at 20 Hz (top right). Opto-sensitive unit histogram during ChR2 excitation at 20 Hz (bottom right). (**D**) No differences in firing rate of optically-sensitive cells between groups (One-way ANOVA $F_{(2, 61)}$=2.703, P=0.0750). (**E**) No differences in burst rate (burst events/s of optically-sensitive cells between groups (One-way ANOVA $F_{(2, 61)}$=0.2806, P=0.7566). (**D-E**, $N_{control}$=25 cells, $N_{Kv4.3LOF}$=13 cells, $N_{BKCa1.1LOF}$=27 cells). Box and whisker data are presented as min to max.

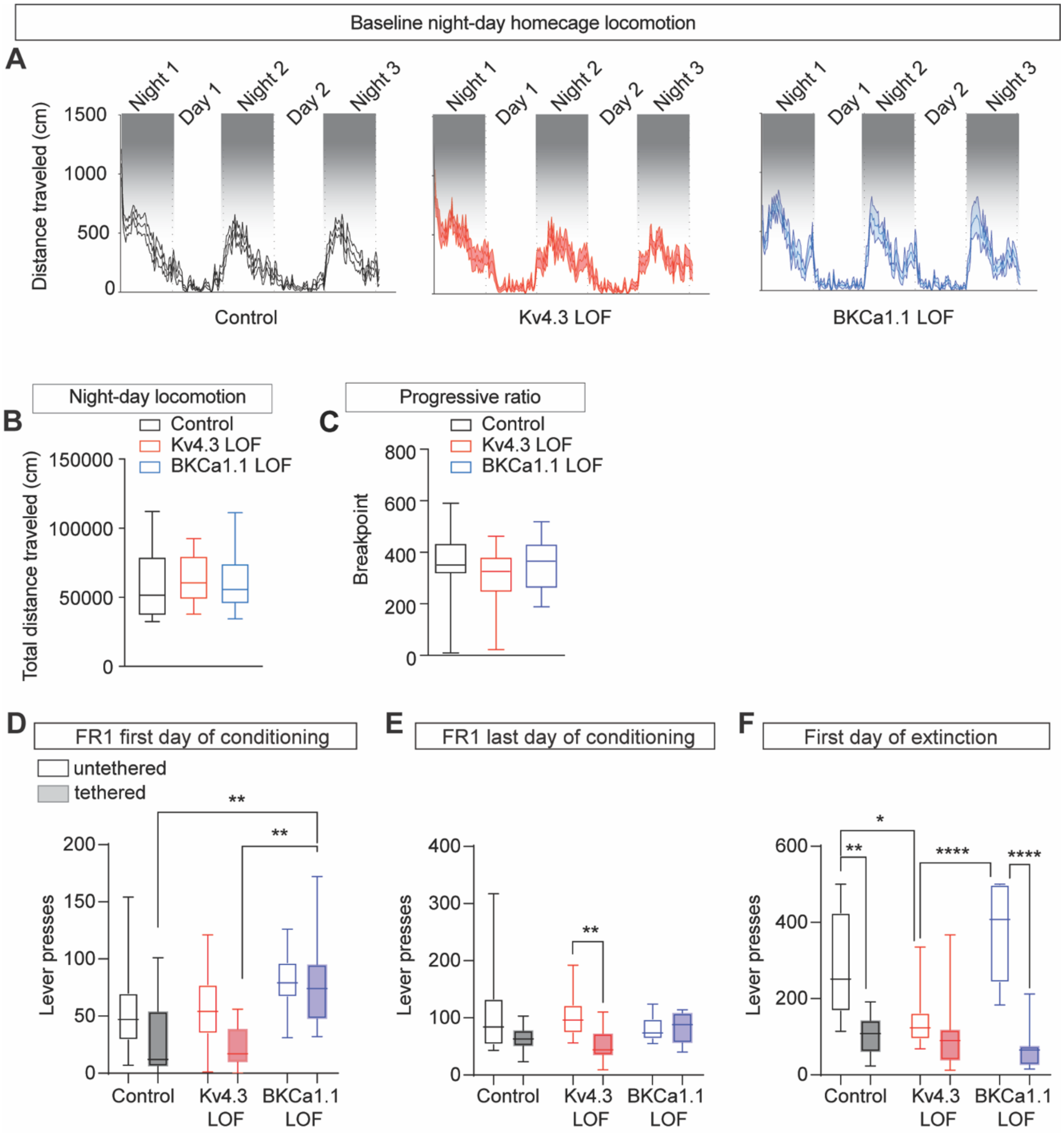


**Fig S6. Additional behavioral analysis of Kv4.3 and BKCa1.1 LOF mice.**

(**A**) Distance travelled in homecage locomotion for control ($N$=15), Kv4.3 LOF ($N$=14), and BKCa1.1 LOF ($N$=12) mice. (**B**) Quantification of total locomotor activity during night and day is not different between groups (One-way ANOVA, F(2, 38)=0.09392, P=0.9106, $N_{control}$=15; $N_{Kv4.3LOF}$=14; $N_{BKCa1.1LOF}$=12). (**C**) Progressive ratio schedule of reinforcement is not different between groups (One-way ANOVA, F(2, 40)=0.7516, P=0.4782, $N_{control}$=14; $N_{Kv4.3LOF}$=16; $N_{BKCa1.1LOF}$=13). (D) Lever press activity for first day of reinforced FR1 conditioning for tethered and untethered mice (Two-way ANOVA, Interaction effect $F_{(2, 90)}$=1.701, P=0.1883; Tethering effect $F_{(1, 90)}$=7.3, P<0.01; Group effect $F_{(2, 90)}$=12.63, P<0.0001; Tukey's multiple comparisons, *p<0.05, **p<0.01, ***p<0.001; untethered $N_{control}$=23, $N_{Kv4.3LOF}$=25, $N_{BKCa1.1LOF}$=18; tethered $N_{control}$=9, $N_{Kv4.3LOF}$= 12,

$N_{BKCa1.1LOF}$=9). (E) Lever press activity for last day of reinforced FR1 conditioning for tethered and untethered mice (Two-way ANOVA, Interaction effect $F_{(2, 89)}$=3.314, P<0.05; Tethering effect $F_{(1, 89)}$=10.04, P<0.01; Group effect $F_{(2, 89)}$=0.207, P=0.8132; Tukey's multiple comparisons, **p<0.01; $N_{control}$=23, $N_{Kv4.3LOF}$=25, $N_{BKCa1.1LOF}$=18; tethered $N_{control}$=8, $N_{Kv4.3LOF}$= 12, $N_{BKCa1.1LOF}$=9). (E) Lever press activity for first day of FR1 extinction conditioning for tethered and untethered mice (Two-way ANOVA, Interaction effect $F_{(2, 59)}$=93.820, P<0.001; Tethering effect $F_{(1, 59)}$=51.39, P<0.0001; Group effect $F_{(2, 59)}$=6.235, P<0.01; Tukey's multiple comparisons, *p<0.05, **p<0.01, ***p<0.001, ****p<0.001; $N_{control}$=13, $N_{Kv4.3LOF}$=10, $N_{BKCa1.1LOF}$=12; tethered $N_{control}$=9, $N_{Kv4.3LOF}$= 12, $N_{BKCa1.1LOF}$=9). Box and whisker data are presented as min to max.

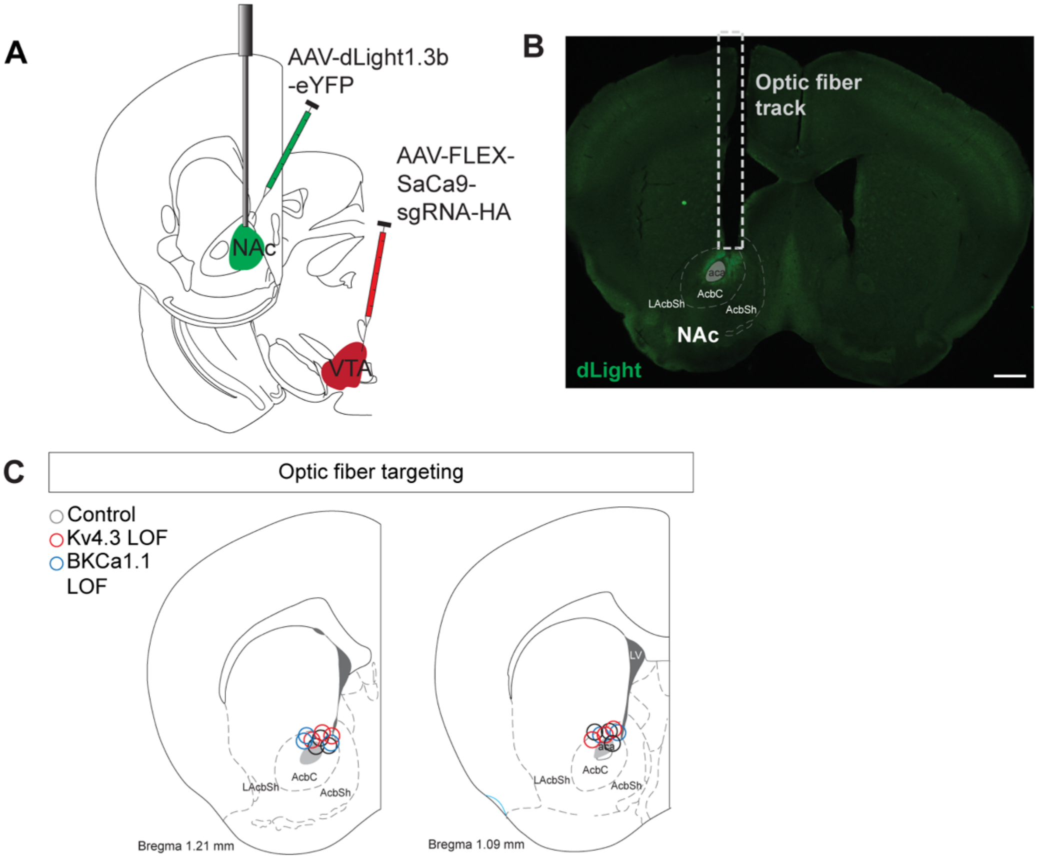


**Fig S7. Targeting of AAV-CAG-dLight1.3 and optic fiber to the NAc.**

(**A**) Schematic of viral injection strategy CRISPR/dLight photometry. (**B**) Representative image of dLight expression in the NAc and optic fiber placement in the NAc. Scale bar= 500 μm. (**C**) Targeting map of all mice.

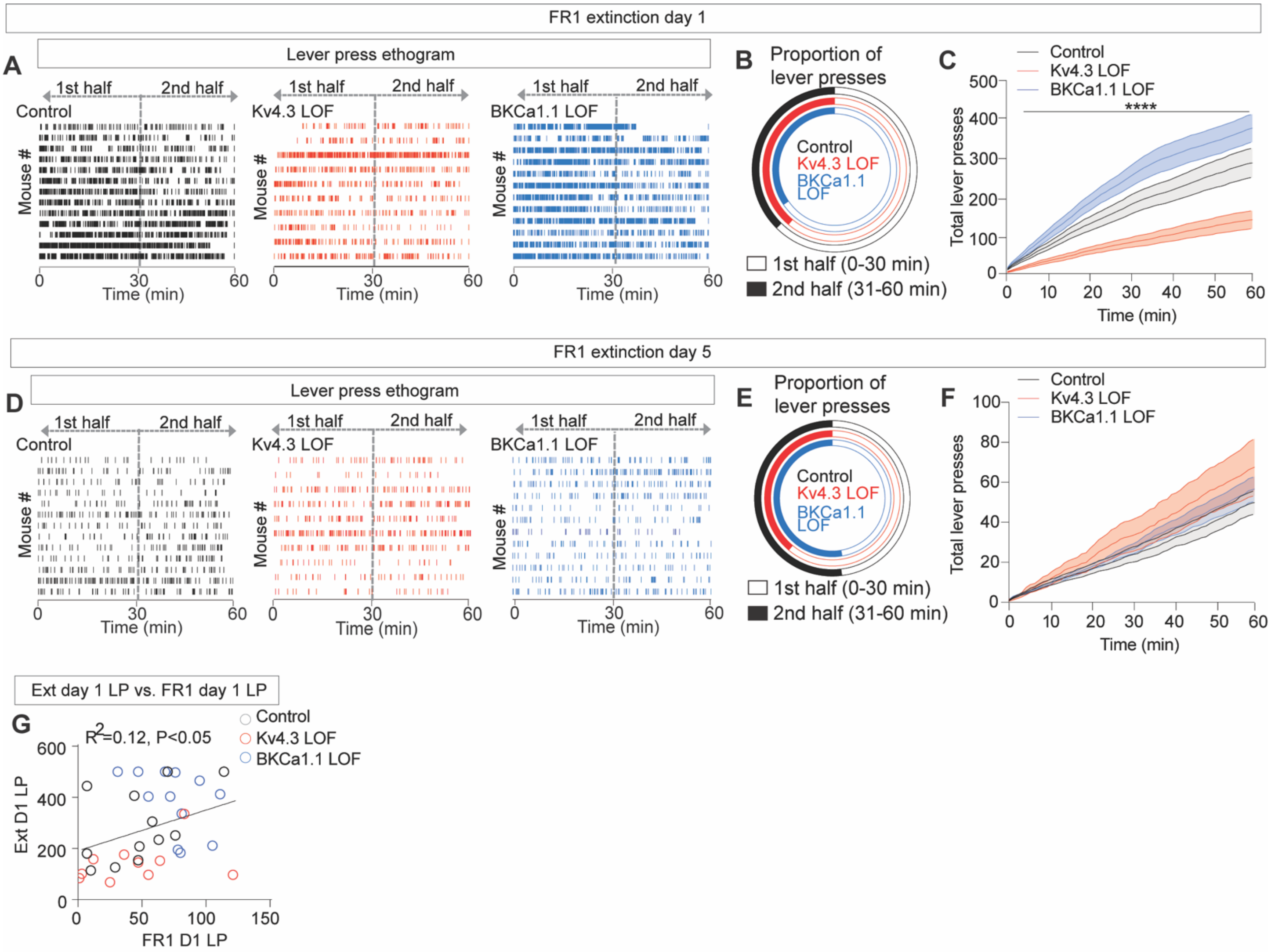


**Fig S8. Extinction learning in Kv4.3 and BKCa1.1 LOF mice.**

(**A**) Lever press ethograms during day 1 of extinction illustrating lower pressing in Kv4.3 LOF group across the session. (**B**) Proportion of lever presses during the first half (0-30 min) and second half (31-60 min) of the day 1 extinction session. (**C**) Cumulative lever presses during day 1 of extinction training (Two-way ANOVA, ****Interaction effect $F_{(118, 1888)}$=8.677, P<0.0001; Time effect $F_{(1.197, 38.30)}$=135.1, P<0.0001; Group effect $F_{(2, 32)}$=12.25, P<0.0001; Subject effect $F_{(32, 1888)}$=207.0, P<0.0001). (**D**) Lever press ethograms during day 5 of extinction illustrating relative uniformity in lever pressing across groups. (**E**) Proportion of lever presses during the first half and second half of the day 5 extinction session. (**F**) Cumulative lever presses during day 5 of extinction training are not different between groups (Two-way ANOVA, ****Interaction effect $F_{(118, 1888)}$=1.205, P=0.712; Time effect $F_{(1.197, 38.30)}$=117.5, P<0.0001; Group effect $F_{(2, 32)}$=1.010, P=0.3754; Subject effect $F_{(32, 1888)}$=169.6, P<0.0001) (**G**) Correlation between total lever presses on day 1 of extinction (Ext D1) and day 1 of FR1 (FR1 D1) (Simple linear regression $F_{(1,32)}$=4.339, *P<0.05). (**A-G**, $N_{control}$=13, $N_{Kv4.3LOF}$=10, and $N_{BKCa1.1}$=12). Data in **C** and **F** are represented as mean $\pm$ SEM).
